# Significance of the CTP-binding motif for the interactions of *S. coelicolor* ParB with DNA, chromosome segregation, and sporogenic hyphal growth

**DOI:** 10.1101/2024.12.04.625172

**Authors:** J. Szymczak, A. Strzałka, D. Jakimowicz, MJ. Szafran

**Affiliations:** Department of Molecular Microbiology, Faculty of Biotechnology, University of Wrocław, Wrocław, Poland; Department of Cell Biology, University of Oklahoma Health Sciences Center, Oklahoma City, Oklahoma, USA

## Abstract

The segregation of bacterial chromosomes is widely mediated by partitioning proteins (ParAB). While ParB binds DNA specifically by recognising short, palindromic sequences known as *parS* sites, ParA utilises its ATPase activity to generate the force to translocate ParB-DNA nucleoprotein complexes (segrosomes). The assembly of the segrosome requires the association of ParB with *parS,* followed by nonspecific spread of the protein along the DNA. To spread on DNA, the ParB dimer must entrap the *parS* site within the complex, a process triggered by CTP binding to the conserved GERR amino acid motif. In *Streptomyces*, a genus of soil-dwelling, multigenomic bacteria that have a complex life cycle, ParB-dependent chromosome partitioning is initiated during the growth of sporogenic hyphae. However, the molecular mechanisms underlying segrosome formation in *Streptomyces* and their ability to coordinate with sporogenic development remain incompletely understood.

In this study, we advance the understanding of chromosome segregation in bacteria by exploring the effects of CTP binding and hydrolysis on the formation of the partitioning complex in *S. coelicolor*. Here, via *in vitro* approaches, we demonstrate that a conserved GERR motif is essential for CTP binding and hydrolysis by *S. coelicolor* ParB. Moreover, the motif is crucial for CTP-dependent ParB accumulation on DNA. Using mutant strains, we show the significance of the GERR motif for segrosome complex assembly. Additionally, we provide data showing that the CTP-binding motif contributes to the regulation of the growth of sporogenic cells. Overall, we show that CTP-dependent segrosome assembly impacts the development of *S. coelicolor* sporogenic cells.

## INTRODUCTION

The faithful segregation of genetic material is a fundamental process of the cell cycle and is a prerequisite for cell propagation (1). In most bacterial species, the segregation of chromosomes and low-copy plasmids is facilitated by a three-component system (ParABS) composed of ParA (*<u>par</u>titioning protein <u>A</u>*) and ParB (*<u>par</u>titioning protein <u>B</u>*) homologues, as well as palindromic 16-base pair (bp) *parS* (*<u>par</u>tition <u>s</u>ite*) sequences (2–4). Depending on the species, one or several *parS* sequences are located near the *oriC* site (*<u>ori</u>gin of <u>c</u>hromosome replicat*i*on*) (5). The *parS* sites are recognised and bound by the ParB protein to form a large, higher-order nucleoprotein complex known as the segrosome (6–8). Essential for specific ParB-*parS* interactions is a centrally positioned DNA-binding domain (DBD), which contains a conserved helix-turn-helix (HTH) motif. The DBD is flanked by the N-(NTD) and C-terminal (CTD) domains, which are required for ParB dimerisation and, through a conserved GLGxG motif, for ParA interactions, respectively (2–4, 9). Additionally, the NTD has recently been demonstrated to be essential for the binding and hydrolysis of cytidine-5’-triphosphate (CTP) via the GERRxR motif, also referred to as the arginine patch (Arg-patch) (10–12).

In the proposed model of bacterial chromosome segregation, the ParB dimer binds the *parS* site (in a process named ParB nucleation) and acts as a CTP molecular switch (12, 13). Upon CTP binding, the conformational changes of the ParB dimer facilitate secondary dimerisation through the NTDs and promote the entrapment of DNA within a clamp-like structure of ParB named the DNA-storing chamber (13, 14). These structural changes subsequently enable the release of the CTP-ParB dimer from the *parS* site and promote the spread across several kilobases along the DNA strand. The DNA-bound ParB can recruit other ParB dimers in a manner independent of the *parS* sequence. This feature allows ParB to cover substantial genomic distance and bypass DNA-bound roadblocks (15). These specific and nonspecific ParB‒DNA interactions lead to DNA bridging and looping, which stimulate DNA compaction. CTP hydrolysis reopens the ParB dimer and leads to cytidine-5’-diphosphate (CDP) removal and dissociation of Apo-ParB from DNA. Interestingly, several *in vitro* and *in vivo* studies have recently suggested that ParB interactions also promote phase separation of ParB-DNA cocondensates (16–18).

Owing to coordinated ParA and ParB activities in most bacteria, one or both segrosomes, formed on the newly replicated *oriC,* are segregated towards the opposite cell poles shortly after *oriC* duplication (2, 10). ParA is an adenosine-5’-triphosphate hydrolase (ATPase) that binds DNA nonspecifically as an ATP‒ParA dimer. The ParB-DNA complex stimulates the ATPase activity of ParA, thereby converting ParA to its monomeric, ADP-bound state, which no longer binds DNA (9, 19, 20). This creates an ATP‒ParA gradient along the chromosome, driving the transport of the CTP‒ParB‒DNA complex along the nucleoid through a ‘diffusion‒ratchet’ mechanism (21, 22).

In *Streptomyces,* chromosome replication and active segregation are temporarily separated due to a unique lifestyle that includes sporulation (23, 24). The *Streptomyces* life cycle starts with spore germination, followed by the apical growth and branching of elongated cells. In contrast to the process in well-studied model species, in *Streptomyces,* chromosomes are extensively replicated during vegetative growth, but cell division rarely occurs. Consequently, the vegetative hyphae are composed of adjacent, elongated cells containing multiple copies of the chromosome, similar to the mycelia of filamentous fungi. Under stress conditions or nutrient limitations, sporogenic cells develop. In these sporogenic cells, chromosomes become actively segregated, aligned along the sporogenic cell compartment, and separated with septa. These events transform multigenomic hyphal cells into chains of spores, each containing a highly compacted copy of the linear chromosome (25).

*Streptomyces* ParB binds numerous *parS* sequences (24 in *Streptomyces coelicolor* (26)) scattered near *oriC*. In the vegetative hyphae of *S. coelicolor*, ParB was shown to form nucleoprotein complexes randomly distributed in cells, except for the apical ParB complex (27–29). Moreover, ParA localises mostly at the apically growing hyphal tip (29). In sporogenic cells, ParA is dispersed throughout the cell and promotes the distribution of segrosomes along the sporogenic hyphae (24). In S*treptomyces,* the ParABS system ensures the precise positioning of multiple chromosomes along sporogenic hyphal cells and synchronises chromosome segregation with cell elongation and multiple divisions (23). Deletion of the *parA* and *parB* genes leads to an increased percentage of anucleate spores and the appearance of minicompartments (23, 29). Moreover, ParA contributes to the regulation of cell extension in *Streptomyces venezuelae* (23, 30). Despite the impact of the ParA and ParB proteins on sporogenic hyphal growth and chromosome partitioning, detailed studies on the significance of CTP for ParB-dependent chromosome segregation in *Streptomyces* have never been conducted.

Here, using a unique, multigenomic model of *S. coelicolor*, we tested how CTP binding affects segrosome assembly. Our findings demonstrated that *S. coelicolor* ParB binds and hydrolyses CTP, which is in line with recent reports on model organisms. We also showed that the GERR motif is crucial for the nonspecific accumulation of the *Sc*ParB protein on DNA. Moreover, we found that the lack of CTP binding leads to the disruption of segrosome complexes in sporogenic *S. coelicolor* cells, affects their growth and results in an increased number of segregation defects.

## MATERIALS AND METHODS

### Bacterial strains and plasmids

The *Escherichia coli* and *S. coelicolor* strains employed in this study are detailed in the Supplementary Materials and Methods (Supplementary Tables 1 and 2). The oligonucleotides and plasmids utilised are specified in the Supplementary Materials and Methods (Supplementary Tables 3 and 4). The culture conditions, antibiotic concentrations, and methods for transformation and conjugation were conducted following standard procedures for *E. coli* (31) and *S. coelicolor* (32). All DNA manipulations were executed in accordance with standard protocols or the manufacturer’s instructions. DNA-modifying enzymes, restriction enzymes, and DNA polymerases were purchased from New England Biolabs (US) or Thermo Fisher Scientific (US). The oligonucleotides were synthesised by Genomed S.A. (Poland), Sigma‒Aldrich (US), or Invitrogen (US).

### Construction of pGEX-6P-1 plasmid derivatives

The pGEX-6P-1 plasmid derivatives encoding *S. coelicolor* ParB (*Sc*ParB) variants, except the pGEX-6P-1_*parB*(HTH) derivative, were constructed on the basis of the pGEX-6P-1_*parB* vector (26). To achieve this, custom synthesis of 325-bp DNA fragments containing specific nucleotide substitutions within the *parB* gene sequence (listed in Supplementary Table 4) was performed (Invitrogen, US). These fragments were digested with the restriction enzymes *PmlI* and *HindIII*. Concurrently, a derivative of pGEX-6P-1_*parB* featuring an 85-bp deletion within the *parB* gene was generated by digestion with *SacI* followed by plasmid religation. The religated plasmid was then digested with *PmlI* and *HindIII,* gel purified, and used for ligation to replace the shortened *parB* gene with the corresponding full-length fragment carrying a point mutation (Supplementary Table 4). To construct the pGEX-6P-1_*parB*(HTH) plasmid, a 935-bp fragment of the *S. coelicolor parB* gene was amplified by PCR using the *Sc*ParB_inside_Fw and ParB_EcoRI_Rv oligonucleotides, with the H24_*parB*_HTH-*SnaBI*-*egfp* cosmid DNA (27) serving as a template. The amplified DNA fragment was digested with the restriction enzymes *PmlI* and *NruI* and subsequently ligated into the *PmlI*- and *NruI*-digested pGEX-6P-1_*parB* plasmid, yielding the pGEX-6P-1_*parB*(HTH) construct. For verification, all the pGEX-6P-1_*parB* derivatives listed in Supplementary Table 4 were tested for the presence of the designated *parB* point mutations by Sanger sequencing.

### Protein overproduction and purification

The pGEX-6P-1_*paB* derivatives were transformed into chemically competent *E. coli* BL21 (DE) pLysS cells. Positive transformants were selected on LB agar plates supplemented with ampicillin and chloramphenicol and subsequently used to overproduce *Sc*ParB variants with N-terminally fused glutathione-S-transferase (GST-*Sc*ParBs). For overproduction, selected transformants were cultured in 800 mL of LB medium supplemented with ampicillin and chloramphenicol. Overproduction of the GST-*Sc*ParB variants was induced by adding 0.5 mM isopropyl β-D-1-thiogalactopyranoside (IPTG) to exponentially growing cells, followed by incubation for 4 hours at 27 °C with shaking at 180 rpm. The cell culture was then centrifuged (20 minutes, 4 °C, 4, 000 rpm), and the supernatant was discarded. The cell pellet was subsequently resuspended in 45 mL of buffer A (100 mM Tris-HCl (pH 8.0), 100 mM NaCl). The cells were sonicated on ice, and the resulting cell lysate was centrifuged (20 minutes, 4 °C, 10, 000 rpm). The clarified lysate was incubated overnight at 4 °C with 1.5 mL of buffer A-equilibrated Glutathione Sepharose® 4B (GE Healthcare, US).

Next, the GST-*Sc*ParB-bound resin was transferred to a plastic column, washed sequentially with 100 mL of buffer A and 50 mL of buffer A supplemented with 450 mM NaCl, and equilibrated with 15 mL of PreScission buffer (100 mM Tris-HCl (pH 8.0), 100 mM NaCl, 10% glycerol, 1 mM 1, 4-dithiothreitol (DTT), and 1 mM ethylenediaminetetraacetic acid (EDTA)). The resin was then incubated overnight at 4 °C with 24 units of PreScission protease (GE Healthcare, US) to cleave the *Sc*ParB variant from the GST tag. The cleaved ScParB variants were eluted, and the fractions with the highest protein concentrations were pooled. The purity of the *Sc*ParB variants was assessed via sodium dodecyl sulfate‒polyacrylamide gel electrophoresis (SDS‒PAGE), followed by staining with InstantBlue^®^ Coomassie protein stain (Abcam, UK). The protein concentration was determined via ROTI^®^-Quant (Carl Roth, US). The purified *Sc*ParB variants were subsequently frozen and stored at -80 °C.

### Analysis of protein‒protein interactions using a bacterial two-hybrid system (BACTH) and β-galactosidase activity assay

To analyse *Sc*ParB dimerisation, *S. coelicolor parB* gene variants were amplified via PCR using the ParB_XbaI_Fw and ParB_KpnI_Rv oligonucleotides, with pGEX-6P-1_*parB* derivatives carrying specific *parB* mutations used as templates (Supplementary Table 4). The PCR products were then purified from an agarose gel, digested with the restriction enzymes *XbaI* and *KpnI*, and ligated into *XbaI-* and *KpnI*-digested pUT18C and pKT25 vectors. The ligation products were transformed into chemically competent *E. coli* DH5α cells, resulting in a collection of pUT18C or pKT25 plasmid derivatives encoding ParB variants (as listed in Supplementary Table 4). All the constructs were verified via Sanger sequencing.

For protein‒protein interaction analysis, the verified pUT18C and pKT25 derivatives encoding specific ParB variants were cotransformed into chemically competent *E. coli* BTH101 cells. The transformants were cultured on LB agar plates supplemented with ampicillin, kanamycin, 0.004% 5-bromo-4-chloro-3-indolyl-β-D-galactopyranoside (X-gal), and 0.5 mM IPTG and incubated for 2 days at 30 °C. Single colonies were subsequently transferred to Eppendorf tubes, resuspended in fresh LB medium, and spotted onto LB agar plates with the same supplements as those described above. After 2 days, the blue pigmentation of the colonies was assessed to verify protein interactions. For the analysis of the interactions of *Sc*ParA and *Sc*ParB, pUT18C derivatives encoding *Sc*ParB variants were cotransformed with the pKT25_*parA* vector into chemically competent *E. coli* BTH101 cells following the procedure described above.

For the β-galactosidase activity assay, selected *E. coli* BTH101 transformants were cultured overnight at 30 °C with shaking at 180 rpm in 3 mL of LB medium supplemented with ampicillin and kanamycin. The next day, 60 µL of the overnight cell cultures was used to inoculate 3 mL of fresh LB medium containing ampicillin, kanamycin, and 0.5 mM IPTG and incubated at 30 °C with shaking at 180 rpm until the optical density (OD) reached approximately 0.4. The cultures were then diluted 5-fold with fresh LB medium, and the OD was measured again. Subsequently, 2.5 mL of the diluted culture was transferred to a fresh 15 mL Falcon tube, and the β-galactosidase activity assay was performed according to a protocol described previously (37). The β-galactosidase activity was quantified in triplicate for each *Sc*ParB variant and expressed in Miller units [Miller U], normalised to the OD of the diluted cell culture. The statistical significance of differences between *Sc*ParB variants was determined using the two-sided Student’s t test.

### Construction of *S. coelicolor* strains

To construct *S. coelicolor* strains producing *Sc*ParB variants with C-terminally fused EGFP (*Sc*ParB-EGFP), we employed the Redirect protocol, which is based on DNA homologous recombination (33). Initially, the *vph-oriT* cassette (viomycin resistance) in the H24_*parB* HTH*-egfp* cosmid (27) was replaced with a *hyg-oriT* cassette (hygromycin resistance) via the Redirect protocol. Next, the cosmid DNA was digested with the restriction enzyme *SnaBI*, purified from an agarose gel, and subsequently recombined with the PCR-amplified fragments of the *parB* gene carrying the point mutations. The *parB* gene fragments were amplified using the ParB_XbaI_Fw and ParB_KpnI_Rv oligonucleotides with the pGEX-6P-1_*parB* derivatives carrying specific nucleotide substitutions within the *parB* gene as the templates (as listed in Supplementary Table 4). The recombination of the linear PCR product and digested cosmid DNA was facilitated by cotransformation into L-arabinose-treated *E. coli* BW25113/pIJ790 cells following the Redirect protocol (33). Positive recombinants were selected on LB agar plates supplemented with hygromycin and verified via colony PCR using the ParB_inside_Fw and ParB_inside_Rv oligonucleotides. Cosmids carrying mutations in the *parB-egfp* gene were then isolated and verified by Sanger sequencing.

The verified cosmids were subsequently used to transform chemically competent *E. coli* ET12567/pUZ8002 cells. The selected transformants were conjugated into the *S. coelicolor* J3303 strain (Δ*parB*) to replace the apramycin resistance gene (*accIV*) in the native *parB* locus with the *parB*-*egfp* variants. The exconjugants were selected on SFM supplemented with kanamycin. To identify double-crossover events, the exconjugants were restreaked on antibiotic-free SFM and screened for sensitivity to apramycin, kanamycin, and hygromycin.

### Electrophoretic mobility shift assay (EMSA)

To analyse the *Sc*ParB-DNA interactions *in vitro*, linear 500-bp DNA fragments containing two *parS* sites were amplified by PCR using the EMSA_*parS*_Fw and EMSA_2x_*parS*_Rv oligonucleotides. For DNA detection, the EMSA_parS_Fw oligonucleotide was custom synthesised with a 5’-conjugated cyanine (Cy) fluorophore, using cyanine 5 (Cy5) for the wild-type *parS*-containing DNA and cyanine 3 (Cy3) for the scrambled *parS-*containing DNA. The DNA templates were pUC19 plasmid derivatives containing either wild-type (pUC19_A7) or scrambled (pUC19_B2) *parS* sequences (36). Following PCR amplification, the 5’-labelled DNA fragments were gel-purified, and 10 µM solutions of the fragments were prepared. These DNA fragments were then mixed with varying concentrations of *Sc*ParB variants (ranging from 200 to 1000) in PreScission buffer in a final volume of 20 µL and incubated for 30 minutes at room temperature. After incubation, 4 µL of 70% glycerol was added to each mixture. The resulting protein‒DNA complexes were resolved via electrophoresis on a 0.8% agarose gel in 1x TBE buffer at a constant voltage of 80 V for 3 hours at 4 °C. Free DNA and DNA‒protein complexes were visualised using an Azure 600 imaging system with filters specific for detecting Cy3 or Cy5 fluorescence. The intensity of Cy5 fluorescence from free DNA was quantified using Fiji software for samples with 500 nM protein. Each experiment was performed in triplicate, and the statistical significance of differences in band fluorescence intensity was evaluated using the two-sided Student’s t test.

### TNP-CTP binding assay

The TNP-CTP-binding assay was performed in a final volume of 200 µL of BLI_A buffer containing 1 µM *Sc*ParB variant and 5 µM 2’, 3’-O-trinitrophenyl-cytidine-5’-triphosphate (TNP-CTP; Jena Bioscience, Germany). Prior to measurement, the mixture was incubated for 30 minutes at room temperature. Optionally, the mixture was supplemented with 34-bp dsDNA containing a *parS* sequence to a final concentration of 2 µM. This DNA fragment contained either the wild-type or scrambled *parS* sequence, obtained by hybridising two complementary 34-bp oligonucleotides: *parS*_34pz_Fw and *parS*_34pz_Rv for wild-type *parS* or mutparS_34pz_Fw and mutparS_34pz_Rv for scrambled *parS*.

Fluorescence spectra were recorded using a Tecan Infinite 200 Pro microplate reader (Tecan Group Ltd., Switzerland) with excitation at 410 nm and emission measured in the range of 400 to 600 nm. The assay was conducted in a 96-well black plastic plate (Black Greiner Bio-One, Thermo Fisher Scientific, US). For the control experiments, *Sc*ParB was subjected to heat inactivation (95 °C for 10 minutes) prior to the addition of TNP-CTP, or wild-type *Sc*ParB was incubated with DNA containing the scrambled *parS* sequence. As a reference, the fluorescence intensity of 5 µM TNP-CTP (or 5 µM TNP-CTP supplemented with *parS*-containing DNA) was measured and set to 1.0 when it was plotted against the fluorescence intensity obtained for *Sc*ParB-containing samples (‘relative fluorescence’). Each experiment was conducted in triplicate, and the statistical significance of the difference between the *Sc*ParB variants was determined using the two-sided Student’s t test.

### CTPase activity assay

The CTPase activity of the ParB variants was quantified using an ATPase/GTPase Activity Assay Kit (Sigma-Aldrich, US). The reaction containing 1 µM ParB variant and 4 mM CTP was performed in BLI_A buffer (100 mM Tris-HCl (pH 8.0), 100 mM NaCl, 1 mM MgCl_2_, 0, 005% Tween 20) in a final volume of 40 µL. Optionally, the reaction was supplemented with 34-bp double-stranded DNA containing the *parS* sequence to a final concentration of 2 µM. The reaction mixture was incubated for 30 minutes at 37 °C. The absorbance at 620 nm was measured in a plastic plate (96-well SpectraPlate, Perkin Elmer, US) using a Tecan Infinite 200 Pro, and used, according to the manufacturer’s description, to calculate the CTP hydrolysis rate as the amount of inorganic phosphate (Pi) released by the *Sc*ParB monomer per hour [Pi/hour]. In the control experiment, *Sc*ParB variants were heat-inactivated by incubating at a temperature of 95 °C for 10 minutes. The measurements for each *Sc*ParB variant were performed in triplicate, and the statistical significance of the difference between ParB variants was determined using the two-sided Student’s t test.

### Circular dichroism (CD) spectroscopy

Circular dichroism (CD) spectra of the *Sc*ParB variants were measured using a J714 CD spectrometer (Jasco, Japan) equipped with a PFD-350S Peltier-type thermostat. Far-UV spectra were recorded at a temperature of 20 °C in a 190 to 250 nm wavelength range using 50 µL of the ParB variant in PreScission buffer and a quartz cuvette with a path length of 10 mm. Each CD spectrum recording was performed in triplicate, and the average ellipticity value was normalised against the molar protein concentration (molar ellipticity [deg * cm^2^ *dmol^−1^]) and plotted versus the wavelength [nm].

### Epifluorescence and structured illumination microscopy (SIM)

For the microscopic analysis of chromosome segregation, *S. coelicolor* cultures were prepared on glass coverslips and stained with DAPI and wheat germ agglutinin (WGA) conjugated to Texas Red (WGA-TexasRed) following a protocol described previously (29), with the exception that cell fixation was carried out by washing the glass-attached mycelia three times with absolute methyl alcohol. Standard epifluorescence observations of the nucleoid (DAPI-stained), cell wall (WGA-TexasRed-stained) or *Sc*ParB-EGFP variants were conducted using a Leica LAS X Widefield microscope equipped with a Fluotar 100x/1.32 objective. The exposure times for the respective fluorophores were as follows: DAPI, 40 ms; Texas Red, 200 ms; and EGFP, 200 ms. The length of prespores was measured as the distance between neighbouring septa. The prespore compartments shorter than 0.5 µm were categorised as minicompartments. Chromosome segregation defects were measured as the percentage of prespore compartments lacking DAPI fluorescence. A total of 510 prespore compartments were analysed for each *S. coelicolor* strain. Additionally, to assess the length of sporogenic hyphae, the distance from the hyphal tip to the basal septum was measured in 50 sporogenic hyphae with clearly visible septa.

For structured illumination microscopy (SIM) observations, *S. coelicolor* cultures were prepared on coverslips and fixed as described above, with the exception that, to visualise the cell wall, hyphae were stained with WGA conjugated to AlexaFluor633 (WGA-AF633) diluted to a final concentration of 2.5 µg/mL in PBS. SIM observations were conducted using a Zeiss Elyra 7 microscope 467 equipped with an Alpha Plan-APO 100x/1.46 Oil DIC VIS lens, sCMOS + emCCD camera and Andor EM-CCD 468 camera. Excitation of the EGFP and AF633 fluorophores was performed at wavelengths of 488 nm and 632 nm, with 10% and 5% of the laser power applied, 5000 mW and 500 mW, at 488 nm and 632 nm, respectively. The exposure time was set to 50 ms for both fluorophores. A series of 15-37 cross sections (Z-stacks) were collected, with a 100 nm distance between frames. The data were preprocessed using Lattice SIM image reconstruction algorithm implemented in Zeiss Zen software, followed by bioinformatics analyses with ImageJ software equipped with Fiji 3D Image Suite (38).

ParB-EGFP complexes were analysed in 18-19 hyphal images with clearly detectable sporogenic septa, which served as markers for *S. coelicolor* sporogenic development. To quantify the ParB-EGFP *loci*, selected regions of the sporogenic mycelium were segmented using the Fiji 3D Image Suite package to generate a 3D view of the ParB-EGFP complexes. Initially, local fluorescence signal maxima were identified using the *3D* ‘Local Maxima’ function. These maxima were then used as seeds for segmentation with 3D Spot segmentation with a Gaussian model function. The segmentation parameters were set as follows: *threshold* = 10000; *radius* = 50; and *standard deviation* = 1.75. R was used to visualise the volume and fluorescence intensity of the detected complexes.

## RESULTS

### CTP Binding and Hydrolysis are Stimulated by the ScParB-*parS* Interactions

The N-terminal domain (NTD) of *S. coelicolor* ParB (*Sc*ParB), similar to that of other ParB homologues, contains a conserved GERRxR amino acid sequence previously shown to bind CTP (Figure 1A and Supplementary Figure 1). To study the impact of CTP on ParB activity in *S. coelicolor*, we first overproduced and purified a set of *Sc*ParB variants carrying amino acid substitutions within the GERR motif. The modifications included single (*Sc*ParB^G140S^, *Sc*ParB^R142A^, and *Sc*ParB^R143A^) or double amino acid substitutions (*Sc*ParB^G140S;R142A^ and *Sc*ParB^R142A;R143A^). These substitutions did not affect the overall spatial structure of the *Sc*ParB variants, as confirmed by CD spectroscopy (Supplementary Figure 2).

**Figure 1.**
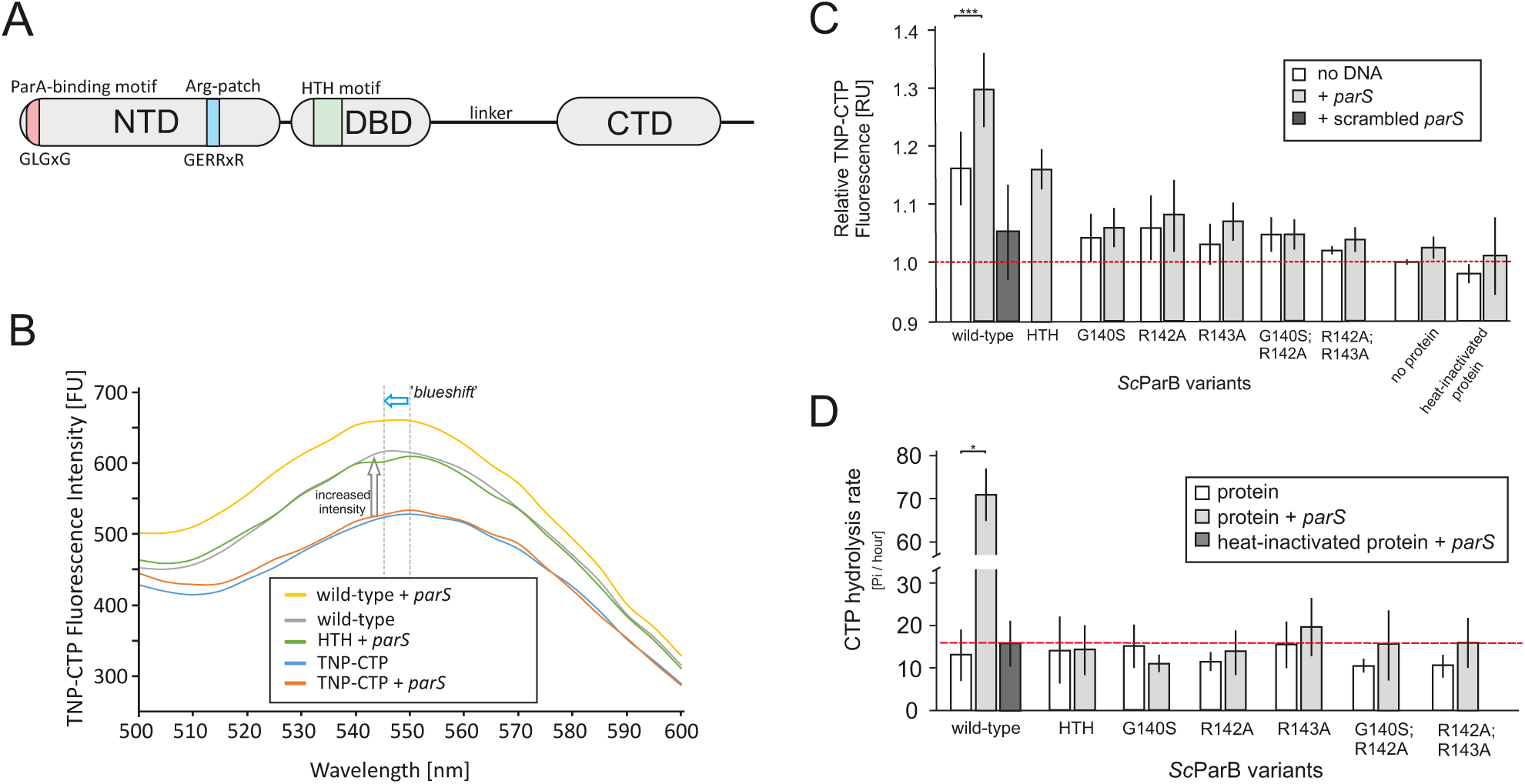
CTP binding and hydrolysis by *Sc*ParB. (**A**) Schematic representation of the subdomain organisation of the *S. coelicolor* ParB protein, highlighting the N-terminal domain (NTD) containing the ParA-binding site (GLGxG), the conserved arginine patch (Arg-patch, GERRxR), the DNA-binding domain (DBD) featuring a helix-turn-helix (HTH) structural motif, and the C-terminal dimerisation domain (CTD), which is connected to the DBD via a flexible linker. **(B)** Representative fluorescence [FU] spectrum of 5 µM TNP-CTP after binding to either wild-type *Sc*ParB or *Sc*ParB^HTH^ (1 µM) in the presence or absence of a 34-bp DNA fragment (2 µM) containing the *parS* sequence. In the control experiments, the fluorescence of 5 µM TNP-CTP and TNP-CTP supplemented with 2 µM 34-bp *parS*-containing DNA was measured. The grey dotted lines indicate the maxima of the fluorescence spectra. The arrows denote the increase in light emission (gray) or the ‘*blueshift*’ in the emission spectrum (blue) following TNP-CTP binding. (**C**) The average relative fluorescence response units [RU] of 5 µM TNP-CTP when incubated with 1 µM *Sc*ParB variants (wild-type, G140S, R142A, R143A, G140S;R142A, R142A;R143A, and HTH) in the absence (white) or presence of the 2 µM 34-bp DNA fragment containing the *parS* (gray) or scrambled *parS* (dark grey) site. The fluorescence of unbound 5 µM TNP-CTP was set to 1.0 and is marked with a red dotted line. (**D**) CTP hydrolysis rate, expressed as µmol of inorganic phosphate (Pi) released per µmol of *Sc*ParB molecules per hour. The analysis was conducted in the presence of 1 µM *Sc*ParB variants (wild-type, G140S, R142A, R143A, G140S;R142A, R142A;R143A, and HTH) and 4 mM CTP and in the absence (white) or presence (gray) of 2 µM 34-bp *parS*-containing DNA. In a control experiment, *Sc*ParB variants were heat-inactivated prior to fluorescence detection (dark grey). The hydrolysis rate of wild-type *Sc*ParB in the absence of *parS*-containing DNA is marked with a red dotted line. All experiments were performed in triplicate, and the quantified standard deviations are indicated. Statistical significance was determined via Student’s t test; p value <0.05 (*), <0.01 (**), and <0.001 (***).

Next, we tested the affinity of the *Sc*ParB variants for a fluorescently labelled CTP analogue, 2’, 3’-O-trinitrophenyl-cytidine-5’-triphosphate (TNP-CTP). The accommodation of TNP-CTP in the binding pocket of *Sc*ParB resulted in an increase in the fluorescence signal (RU, response units) and shifted the maximum emitted fluorescence signal towards a shorter wavelength (*‘blueshift’*) (Figure 1B, and Supplementary Figure 3). The relative fluorescence signal of TNP-CTP increased when the nucleotide was incubated with wild-type *Sc*ParB (1.16 ± 0.06 RU) compared with that in the control experiments with no protein added (1.00 ± 0.00 RU) or when the wild-type *Sc*ParB was first heat inactivated (0.98 ± 0.02 RU) (Figure 1C). For all the GERR-substituted *Sc*ParB variants, the detected TNP-CTP fluorescence (from 1.02 to 1.07 RU) was comparable to that of the negative controls (Figure 1C), indicating that an intact GERR motif is required for CTP binding.

Having established that *Sc*ParB binds CTP, the next step was to investigate whether the formation of the ScParB-*parS* complex affects the affinity of *Sc*ParB for the nucleotide. To this end, we incubated the wild-type *Sc*ParB or GERR-substituted variants with TNP-CTP and a 34-bp long DNA fragment containing a single *parS* sequence. We observed that the addition of *parS*-containing DNA stimulated TNP-CTP binding. The relative fluorescence signal in the presence of wild-type *Sc*ParB (1.29 ± 0.06 RU) was significantly increased than the signal in the absence of DNA (1.16 ± 0.06 RU) or in the presence of a scrambled *parS* sequence (1.04 ± 0.08 RU) (Figure 1C). As predicted, for all GERR-substituted *Sc*ParB variants, which did not interact with TNP-CTP, the relative fluorescence signals in the presence or absence of DNA were comparable (Figure 1C). The increased affinity of *Sc*ParB for TNP-CTP in the presence of *parS* was not detected for the non-DNA-interacting *Sc*ParB^HTH^ variant. The TNP-CTP relative fluorescence signal (1.16 ± 0.03 RU) for this variant in the presence of *parS* was comparable to that of wild-type *Sc*ParB without the addition of *parS* (1.16 ± 0.06 RU) (Figure 1C). This finding indicates that although *Sc*ParB binds CTP *in vitro* in the absence of *parS*-containing DNA, its affinity for the nucleotide is increased when *Sc*ParB binds *parS*.

Next, we tested whether the presence of the *parS* sequence impacts not only CTP binding but also CTP hydrolysis (CTPase activity). For the wild-type *Sc*ParB protein in the absence of *parS*, the CTP hydrolysis rate was low, with 13.0 ± 4 Pi/hour released per ParB monomer. Since a similar CTP hydrolysis rate was detected for heat-inactivated *Sc*ParB (15.6 ± 6 Pi/hour), we estimated the rate quantified above as spontaneous CTP degradation under the assay conditions. When the reaction mixture was supplemented with *parS*-containing DNA, the CTPase activity of wild-type *Sc*ParB notably increased to 71.1 ± 6 Pi/hour (Figure 1D). As predicted, a lack of CTPase activity (8.4 to 19.4 Pi/hour) was observed for the *Sc*ParB variants with GERR motif substitutions, which is consistent with their lack of CTP binding (Figure 1D). Moreover, the CTPase activity of *Sc*ParB^HTH^, which interacts with CTP but not with DNA (Supplementary Figure 4A), was completely abolished (14.1 ± 6 Pi/hour). These findings confirm that the interaction of *Sc*ParB with *parS* increases its CTPase activity.

Taken together, our observations confirmed that *S. coelicolor* ParB is a CTP-binding protein. CTP binding is facilitated by interactions with glycine and two arginine residues within the conserved GERR motif located in the NTD. Moreover, our findings also indicate that the CTP-*Sc*ParB complex can be formed in the absence of *parS,* but nucleotide binding is stimulated as a result of the ScParB-*parS* interaction. The interaction of the CTP-*Sc*ParB complex with *parS* enhances the CTPase activity of the protein‒nucleotide complex.

### CTP Does Not Contribute to the Association of *Sc*ParB with *parS*

Given that the GERR motif is essential for CTP binding and hydrolysis, we tested whether the absence of CTP binding affects the interactions of *Sc*ParB with DNA. First, via an EMSA, we examined the interactions of *Sc*ParB variants with a 500-bp long, Cy5-labelled DNA fragment containing two *parS* sites (compared with Cy3-labelled DNA with scrambled *parS*). The wild-type *Sc*ParB bound specifically to *parS*-containing DNA at the lowest tested concentration (250 nM) (Figure 2A), discriminating against the scrambled *parS* sites used in the control assay (Supplementary Figure 4A). At 250 nM *Sc*ParB, only a single *Sc*ParB-*parS* complex was detected, whereas at 500 nM, two separate ParB-*parS* complexes were observed, and the amount of free DNA remained below 10%. Since no large protein complex was detected, this observation suggests that the association of *Sc*ParB with both *parS* sequences (*Sc*ParB nucleation) may occur independently (Figure 2A). Additionally, the presence of 1 mM CTP in the reaction mixture did not affect the assembly of the protein‒DNA complex, suggesting that CTP has no effect on *Sc*ParB nucleation on *parS* sequences (Supplementary Figure 4B). All the tested *Sc*ParB variants interacted with *parS*-containing DNA with a similar affinity to that of the wild-type *Sc*ParB, except for the *Sc*ParB^G140S;R142A^ variant, the DNA binding of which was diminished, resulting in a greater fraction of unbound DNA at a 500 nM protein concentration (Figure 2A). As expected, the presence of CTP did not affect the *parS* binding of the GERR-substituted *Sc*ParB variants (Supplementary Figure 4C).

**Figure 2.**
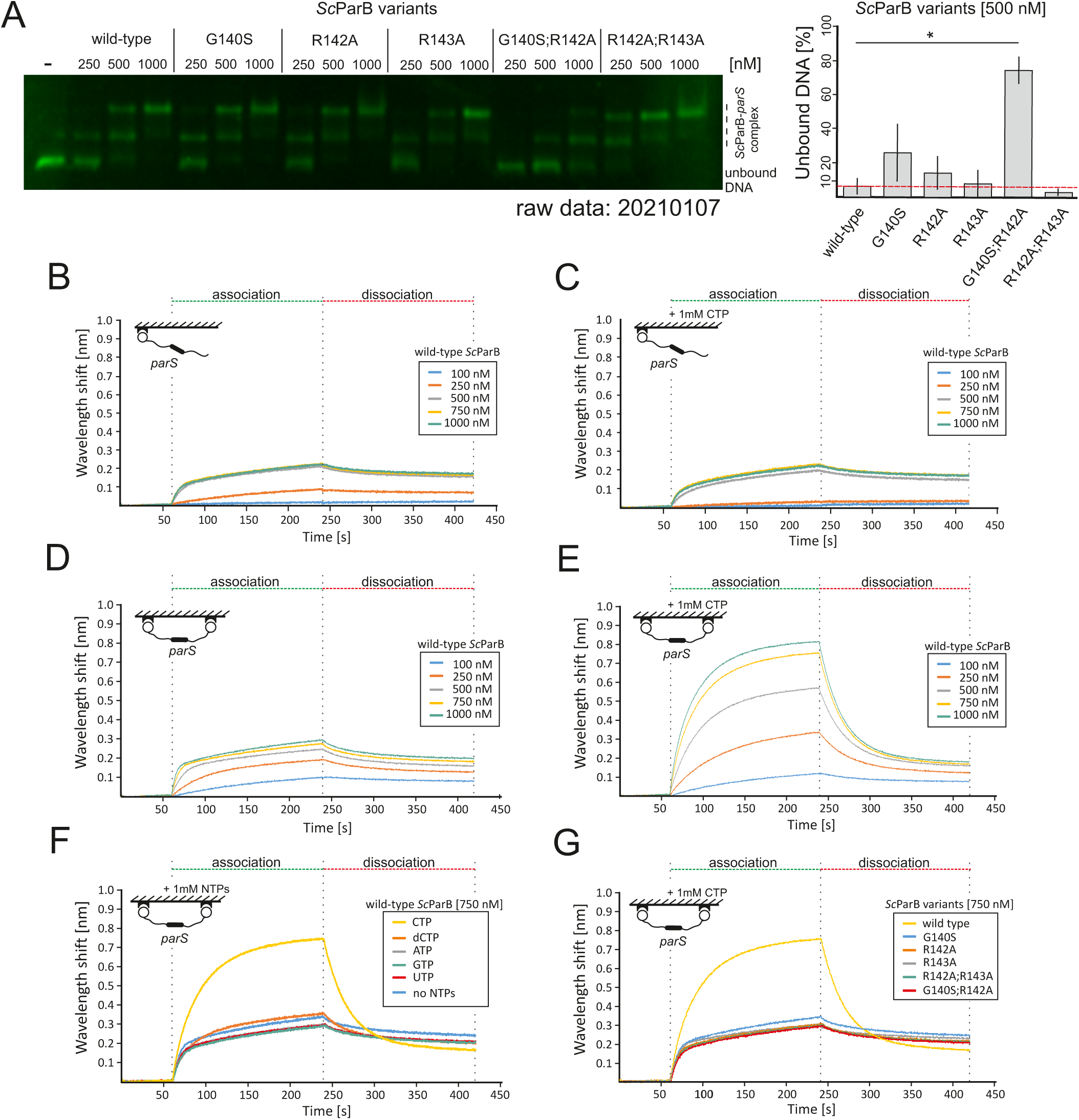
Impact of CTP on *Sc*ParB-DNA interactions *in vitro.* (**A**) *Left*: Analysis of the binding of recombinant *Sc*ParB variants (wild-type, G140S, R142A, R143A, G140S, R142A, and R142A;R143A) to a 500-bp Cy5-labelled linear DNA fragment (10 µM) containing two *parS* sites. The *Sc*ParB concentrations ranged from 250 to 1000 nM. *Right*: The unbound DNA fraction (%) was quantified with a *Sc*ParB variant concentration of 500 nM. The experiment was conducted in triplicate, with the quantified standard deviations indicated. Statistical significance was determined via Student’s t test; p values <0.05 (*), <0.01 (**), and <0.001 (***). (**B**) BLI-measured binding of wild-type *Sc*ParB (100–1000 nM protein) to a one-end biotin-immobilised 300-bp DNA fragment containing a wild-type *parS* site, also (**C**) conducted in the presence of 1 mM CTP. (**D**) BLI-measured binding of wild-type *Sc*ParB (100–1000 nM protein) to a one-end biotin-immobilised 300-bp DNA fragment containing a wild-type *parS* site, also (**E**) conducted in the presence of 1 mM CTP. (**F**) BLI-measured binding of 750 nM wild-type *Sc*ParB to a two-end biotin-immobilised 300-bp DNA fragment containing a wild-type *parS* site performed in the presence of various nucleotide-5’-triphosphates (CTP, dCTP, ATP, GTP, or UTP) at 1 mM. (**G**) BLI-measured binding of 750 nM *Sc*ParB variants (wild-type, G140S, R142A, R143A, G140S;R142A, and R142A;R143A) to a two-end biotin-immobilised 300-bp DNA fragment containing a wild-type *parS* site performed in the presence of 1 mM CTP. The association and dissociation steps are indicated by green and red dotted lines, respectively.

The results of the EMSAs were further confirmed by detailed analysis of the *Sc*ParB-DNA interactions using BLI. In this experiment, we used a 300-bp *parS*-containing DNA fragment biotinylated at the 5’-end and immobilised on the BLI sensor surface (Figure 2B, inset). The binding of wild-type *Sc*ParB to DNA was observed at a protein concentration of 250 nM, whereas at a concentration of 500 nM (or higher), the detected signals were comparable and reached a plateau (Figure 2B). The dissociation constant (K_d_) of the ScParB-*parS* complex for one-end immobilised DNA was 271 ± 14 nM (Supplementary Figure 5). The addition of 1 mM CTP to the association buffer did not significantly affect the association curve, and the saturation of *parS* sites was achieved at protein concentrations comparable to those under CTP-free conditions (Figure 2C). This observation reinforces the EMSA result indicating that CTP is not required for *Sc*ParB-*parS* binding.

### CTP Binding is Required for *Sc*ParB Accumulation on DNA

Given that no high-molecular-weight protein‒DNA complex was detected in our EMSA or BLI studies, we further investigated the affinity of *Sc*ParB for the DNA substrate that enables the spread of ParB. To this end, as in the above-described experiment, a 300-bp long DNA fragment was attached to the surface at both ends, creating two spatial ‘*roadblocks*’ with a single *parS* site located between them (Figure 2D, inset). BLI analyses revealed that in the absence of CTP, *Sc*ParB bound to *parS*-containing DNA attached at two ends (Figure 2D) with an affinity comparable (167 ± 27 nM, Supplementary Figure 5) to that observed for DNA with one end immobilised (Figure 2B, and Supplementary Figure 5). Next, we compared the interaction of *Sc*ParB with *parS*-containing two-end immobilised DNA in the presence of 1 mM CTP (Figure 2E). The addition of 1 mM CTP strongly stimulated *Sc*ParB binding to DNA with both ends immobilised, which suggested the accumulation of the protein on the DNA. The CTP-dependent accumulation of *Sc*ParB on DNA was observed even at the lowest tested CTP concentration (63 µM) and increased at higher CTP concentrations (Supplementary Figure 6A). Further analysis confirmed that *Sc*ParB accumulation on DNA occurs specifically only in the presence of the *parS* site, as it was not detected for the *Sc*ParB^HTH^ variant or on DNA with a scrambled *parS* site (Supplementary Figures 6B-D). The accumulation of *Sc*ParB on DNA was also not observed in the presence of other nucleotide-5’-triphosphates (ATP, GTP, or UTP) or deoxycytidine-5’-triphosphate (dCTP), indicating the strong specificity of the CTP-binding pocket for the coordinated nucleotide (Figure 2F). The impact of CTP on *Sc*ParB-DNA interactions was further verified by analysing *Sc*ParB variants with GERR motif substitutions. Although these *Sc*ParB variants interacted with *parS*-containing DNA, they were unable to accumulate on DNA in the presence of CTP (Figure 2G), confirming the significance of CTP binding for *Sc*ParB accumulation on DNA.

In summary, our observations reinforce the model proposed for ParB homologues in other bacterial species, which posits that nonspecific *Sc*ParB accumulation on DNA requires both the presence of *parS* sequences and CTP binding. We infer that the observed accumulation reflects *Sc*ParB spreading from the *parS* site.

### Lack of CTP Binding by *Sc*ParB Affects Segrosome Assembly in *S. coelicolor*

Given that CTP binding promotes *Sc*ParB accumulation on DNA, we next investigated how the elimination of CTP binding influences segrosome formation in *S. coelicolor* sporogenic hyphae, i.e., at the stage of segregation of multiple chromosomes into unigenomic prespores. For this purpose, we constructed *S. coelicolor* strains producing GERR-substituted *Sc*ParB variants, each C-terminally fused with EGFP (*Sc*ParB^G140S^-EGFP, *Sc*ParB^R142A^-EGFP, and *Sc*ParB^G140S;R142A^-EGFP). These strains were compared to those producing the wild-type *Sc*ParB-EGFP or its non-DNA-interacting *Sc*ParB^HTH^-EGFP variant (Supplementary Figure 7). Despite numerous attempts, we were not successful in the selection of *S. coelicolor* strains producing R143A-substituted *Sc*ParB variants, namely, *Sc*ParB^R143A^-EGFP and *Sc*ParB^R142A;R143A^-EGFP. First, we examined the formation of segrosomes in *S. coelicolor* strains producing *Sc*ParB-EGFP variants (Figure 3A). In the strain producing the wild-type *Sc*ParB-EGFP variant, bright and uniformly spaced *Sc*ParB-EGFP *foci* were observed between the septa, forming chains of prespores, as described previously (27, 39). In contrast, the EGFP fluorescence in the strains producing *Sc*ParB variants deficient in CTP binding, namely, *Sc*ParB^G140S^-EGFP and *Sc*ParB^G140S;R142A^-EGFP, was highly diffuse along the sporogenic hyphae, similar to the fluorescence signal observed in the strain producing *Sc*ParB^HTH^-EGFP, in which DNA binding was abolished.

**Figure 3.**
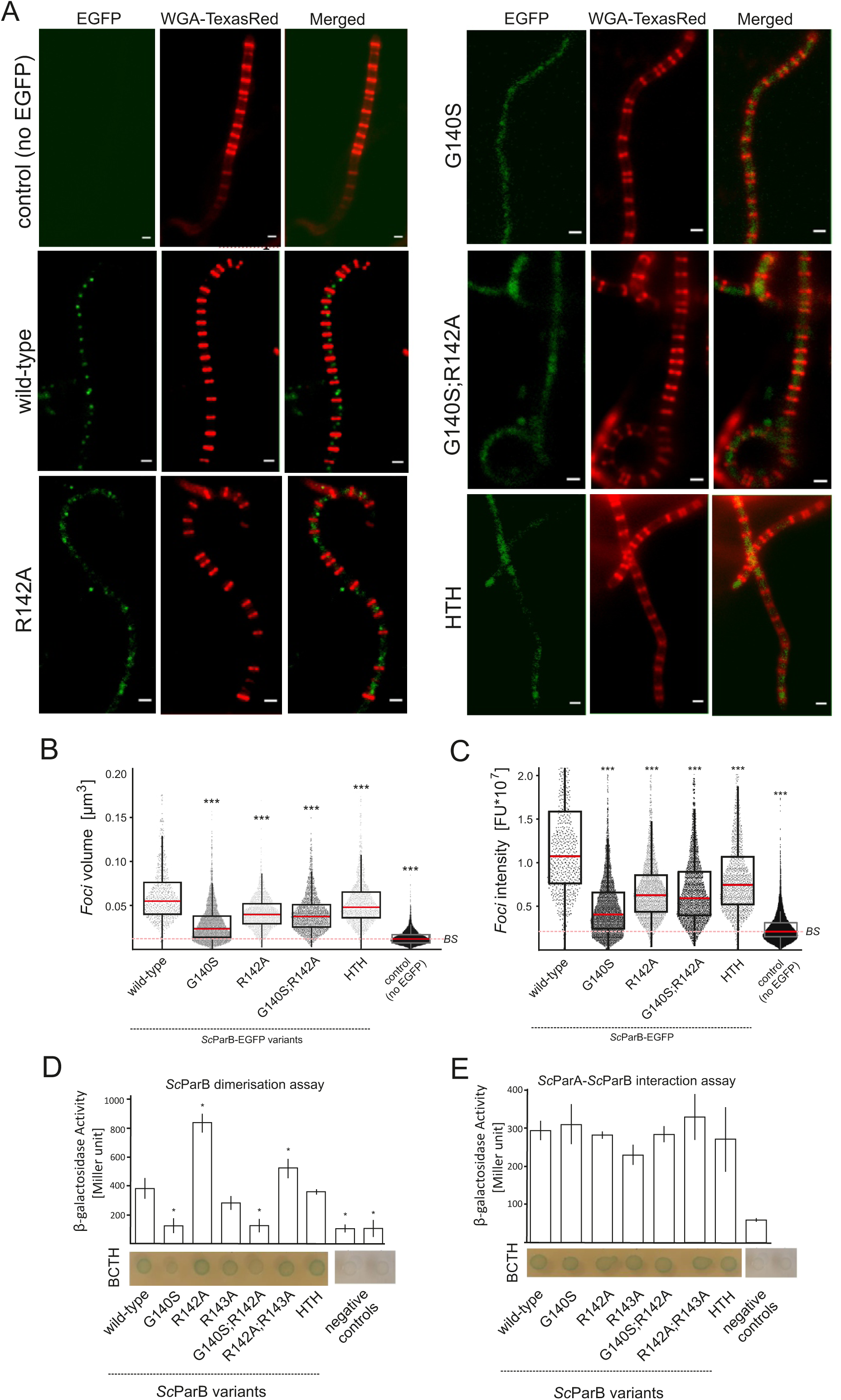
Effect of GERR motif substitution on *Sc*ParB-EGFP complex assembly. (**A**) Visualisation of fluorescence *foci* in *S. coelicolor* sporogenic hyphae of strains producing *Sc*ParB-EGFP (wild-type, G140S, R142AP, G140S;R142A, and HTH). The peptidoglycan was visualised using wheat germ agglutinin conjugated to Texas Red (WGA-TexasRed). The wild-type *S. coelicolor* strain, which does not produce EGFP-fused *Sc*ParB, served as a control. Scale bar: 2 µm. (**B**) Dimerisation capacity or (**C**) *Sc*ParA interaction of *Sc*ParB variants (wild-type, G140S, R142A, R143A, G140S;R142A, R142A;R143A, and HTH) fused with T25 or T18 adenylate cyclase subunits. For the *Sc*ParA-*Sc*PArB interaction assay, *Sc*ParA was fused with the T18 subunit, whereas *Sc*ParB was fused with the T25 subunit. Protein interactions were quantified using a bacterial two-hybrid (BACTH) system or β-galactosidase activity assay [with results expressed in Miller units]. The negative controls included *E. coli* BTH101 cells cotransformed with the pKT25 plasmid and wild-type *parA*- or *parB*-expressing plasmids (pUT18C-*parA* or pUT18C-*parB,* respectively) or the pKT25 and pUT18 plasmids. Box plots representing the volume (**D**) and intensity (**E**) of fluorescence *foci* in strains producing *Sc*ParB-EGFP variants, as identified using SIM. Each boxplot shows the median (red) with the first and third quartiles, whereas the lower and upper ‘whiskers’ extend to values no further than 1.5 times the interquartile range. Statistical significance was determined via Student’s t test; p values <0.05 (*), <0.01 (**), and <0.001 (***). The baseline signal (BS) is indicated with a red dotted line.

High-resolution SIM was subsequently employed to further characterise the *Sc*ParB-EGFP *foci*. SIM analyses allowed measurement of both the intensity and volume of *Sc*ParB-EGFP complexes in the studied *S. coelicolor* strains. The average volume of the wild-type *Sc*ParB-EGFP complexes was 0.062 µm^3^, with a mean intensity of 1.2 × 10^7^ fluorescence units (FU), which was significantly greater than that detected in the other strains. In contrast, the *Sc*ParB *foci* detected in the strain producing the ParB^G140S^-EGFP variant had both the smallest volume (0.029 µm^3^) and the lowest fluorescence intensity (0.4 × 10^7^ FU). However, other GERR-substituted ParB-EGFP variants also presented *foci* with low intensity (Figure 3D) and volume (Figure 3E). The SIM analyses also revealed small *foci* in the strain producing *Sc*ParB^HTH^-EGFP, with an average *focus* volume of 0.054 µm^3^ and an intensity of 0.8 × 10^7^ FU. Although both the average *foc*us volume and intensity of *Sc*ParB^HTH^-EGFP were significantly lower than those detected for the wild-type *Sc*ParB-EGFP, the detected signals were above the BS threshold. Intriguingly, their volume and intensity were slightly greater than those of the GERR-substituted *Sc*ParB-EGFP variants. Notably, autofluorescence was also detected in the wild-type *S. coelicolor* M145 strain, which does not produce the EGFP-fused ScParB protein (average *focus* volume of 0.012 µm³ and intensity of 0.2 × 10⁷ FU (‘*background signal*’; *BS*)). However, the autofluorescence was significantly lower than that in *S. coelicolor* strains producing EGFP-fused *Sc*ParB variants (Figure 3D and 3E). Thus, SIM analyses confirmed that the GERR substitutions reduced the ability of ParB-EGFP to form *foci*. Surprisingly, when we verified the impact of the G140S substitution on the *Sc*ParB-ScParB interaction using the bacterial two-hybrid system and β-galactosidase activity assay, we noticed that the *Sc*ParB^G140S^ and *Sc*ParB^G140S;R142A^ variants were also defective in ParB dimerisation (Figure 3B) but not in the interaction with *Sc*ParA (Figure 3C). On the other hand, the R142A substitution had the opposite effect and resulted in 2-fold enhancement of the association of the *Sc*ParB^R142A^ variant than with that of wild-type *Sc*ParB (Figure 3B). Intriguingly, the increased dimerisation of the R142A-substituted variant corroborated the observation that the *S. coelicolor* strain producing the *Sc*ParB^R142A^-EGFP variant still exhibited detectable fluorescent *foci,* although these *foci* were nonuniformly intense and unevenly positioned compared with those of the wild-type *Sc*ParB-EGFP (Figure 3A).

In summary, using EGFP-tagged *Sc*ParB variants, we demonstrated that the lack of CTP binding significantly affects segrosome assembly during sporulation of *S. coelicolor*. Substitutions within the GERR motif result in less efficient assembly of the *Sc*ParB-DNA complexes and/or their misplacement in sporogenic cells. Moreover, we showed that modifications within the GERR motif can also impact the ability of *Sc*ParB to dimerise but not to interact with *Sc*ParA.

### Substitutions within the GERR Motif Affect *S. coelicolor* Sporulation

Previous studies have shown that *parB* deletion in *S. coelicolor* increases the percentage of anucleate spores and results in irregularly laid septa, leading to the formation of minicompartments (40). Here, we revealed that the formation of *Sc*ParB-DNA complexes *in vitro* and *in vivo* was strongly affected by the elimination of CTP binding caused by GERR motif substitutions. Therefore, to further assess the phenotypic effects of the introduced mutations, we analysed their impact on chromosome segregation and septation in *S. coelicolor* sporogenic hyphae. To this end, we used a set of previously characterised *S. coelicolor* strains producing wild-type *Sc*ParB and GERR-substituted *Sc*ParB variants fused with EGFP. These strains were stained with DAPI to visualise the nucleoids and with WGA-TexasRed to detect sporogenic septa (Figure 4A). Detailed analysis of prespore length revealed that, compared with the wild-type ScParB-EGFP-producing strain (or the control strain with no EGFP fusion), only the *S. coelicolor* strains producing *Sc*ParB variants with the R142A substitution (including the double R140S;R142A substitution) presented an aberrant distance between septa and an increased number of minicompartments (prespore length less than 0.5 µm) (Figure 4B). Moreover, the introduced mutations led to the formation of prespore compartments lacking nucleoids. The percentage of anucleate prespores was 2.7% and 3.4% in both control strains producing the wild-type *Sc*ParB protein and those producing the EGFP-fused variant, respectively. In the strains producing the *Sc*ParB^G140S^-EGFP or non-DNA-interacting *Sc*ParB^HTH^-EGFP variant, the number of anucleate prespores was elevated to 7.6%. Intriguingly, the production of the *Sc*ParB^G140S;R142^-EGFP variant increased the number of prespores lacking DNA to 12.5%, which was greater than that in the strain producing the non-DNA-interacting *Sc*ParB^HTH^-EGFP variant (Figure 4B).

**Figure 4.**
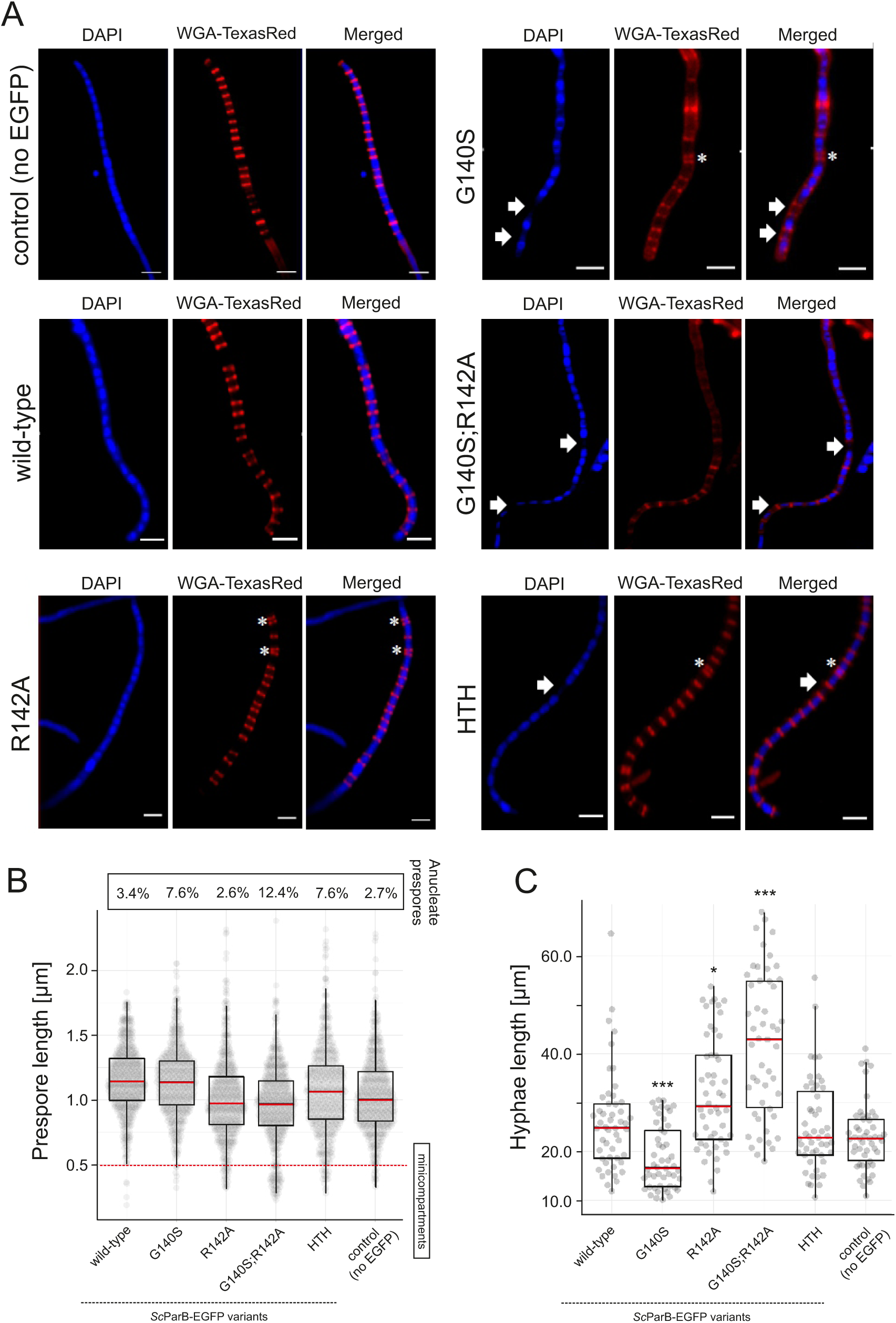
Effect of GERR substitutions in *Sc*ParB-EGFP on *S. coelicolor* chromosome segregation and prespore formation. (**A**) Localisation of nucleoids (DAPI-stained) and peptidoglycan (stained with wheat germ agglutinin, WGA-TexasRed) in *S. coelicolor* strains producing *Sc*ParB-EGFP variants (wild-type, G140S, R142A, G140S;R142A and HTH). The wild-type *S. coelicolor* strain, which does not produce the EGFP-fused *Sc*ParB protein, served as a control. Asterisks indicate the positions of the minicompartments (<0.5 µm), and arrows highlight prespores lacking DAPI fluorescence. Scale bar: 2 µm. (**B**) Box plots displaying the distribution of prespore lengths (n = 510 for each strain). The red dotted lines indicate minicompartments. The percentage of prespore compartments lacking a DAPI signal is shown above. (**C**) Box plot analysis of the lengths of sporogenic hyphae in *S. coelicolor* strains producing *Sc*ParB-EGFP variants (wild-type, G140S, R142A, G140S;R142A, and HTH). The wild-type *S. coelicolor* strain, which does not produce EGFP-fused *Sc*ParB, served as a control. The length from the hyphal tip to the farthest detectable septum was measured on the basis of 50 images of sporogenic hyphae collected from each strain. All box plots show the median (red) with the first and third quartiles, whereas the lower and upper ‘whiskers’ extend to values no further than 1.5 times the interquartile range.

Next, since ParA and ParB were shown to affect the extension of sporogenic cells, we analysed the length of sporogenic cells in the constructed mutant strains. The production of the G140S-substituted variant resulted in the formation of significantly shorter sporogenic hyphae (18.7 ± 6.4 µm) than those produced by the strains producing the wild-type *Sc*ParB (with or without EGFP fusion, 26.1 ± 10.1 µm and 22.6 ± 7.4 µm, respectively) or *Sc*ParB^HTH^ (25.2 ± 9.8 µm) (Figure 4C). On the other hand, in the strain producing *Sc*ParB^R142A,^ the length of sporogenic hyphae increased to 32.0 ± 11.4 µm, whereas this increase was even more pronounced in the strain producing *Sc*ParB^G140S;R142A^ (42.1 ± 14.6 μm). Interestingly, the elongated sporogenic hyphae detected in the strain producing *Sc*ParB^G140S;R142A^ corroborate the observation that the growth of the strain in liquid medium was also accelerated compared with that of the wild-type strain (or strains with single GERR motif substitutions) (Supplementary Figure 8).

In summary, our results showed that abolishing of CTP binding by *Sc*ParB affects the development of sporogenic cells and disturbs chromosome segregation.

## DISSCUSION

Here, we showed that the *S. coelicolor* partitioning protein ParB binds and hydrolyses CTP in a *parS*-dependent manner. Moreover, CTP binding, which is mediated by the GERR motif located within the NTD, is essential for *Sc*ParB accumulation on DNA and segrosome complex formation.

In all the proposed models, the first step of segrosome assembly is the specific binding of ParB to the *parS* site (4, 13, 41)*. S. coelicolor* ParB has been reported to form a complex with *parS* with a dissociation constant (K_d_) that varies from 33 nM (42) to 480 nM (26). However, our findings did not corroborate these observations, as the BLI-estimated K_d_ values ranged from 176 nM to 271 nM, depending on the immobilised DNA. We demonstrated that in *S. coelicolor,* both Apo-*Sc*ParB and CTP-*Sc*ParB exhibit similar affinities for the *parS* site, suggesting that the nucleation step is CTP independent (Figure 5, A-D). To our knowledge, the intracellular CTP levels in *S. coelicolor* have not been determined. However, the CTP concentration in other bacteria, which is tenfold greater than the K_d_ of the CTP-ParB complex (∼ 10 µM), suggests preloading of *Sc*ParB with CTP (CTP-*Sc*ParB) before it binds to *parS* (11). Intriguingly, CTP binding to *C. crescentus* ParB (*Cc*ParB) was not detected in the absence of parS-containing DNA or was found to be very low for *B. subtilis* (*Bs*ParB) (10, 11).

**Figure 5.**
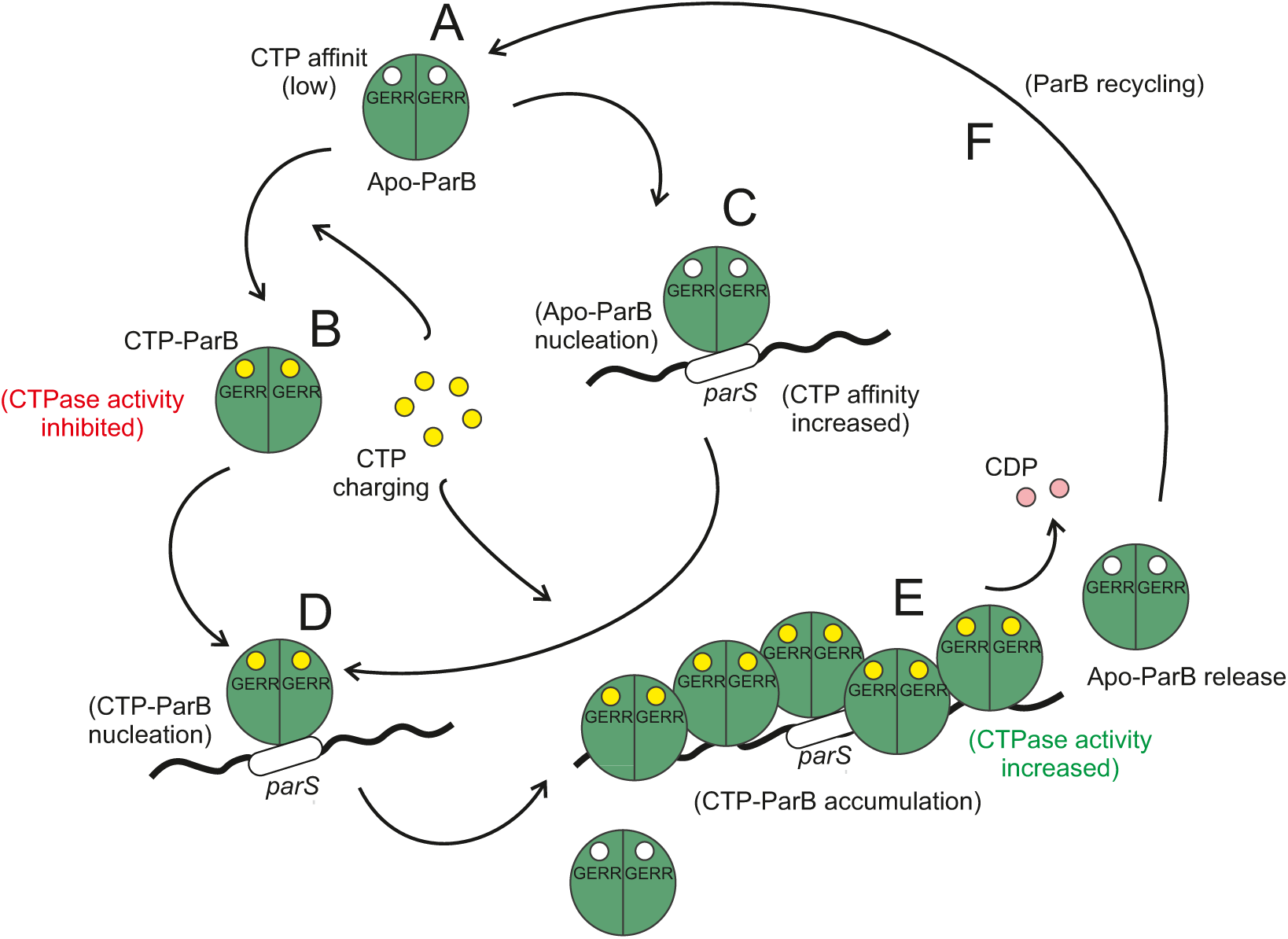
Model of *Sc*ParB interactions with DNA.

We speculate that, at least in *Streptomyces*, the initial step of *Sc*ParB nucleation is independent of nucleotide availability, as all tested GERR-substituted variants bind *parS*-containing DNA with affinity comparable to that of the wild-type protein. Our observations are also supported by studies on the *C. crescentus* ParB (*Cc*ParB) homologue, in which the *Cc*ParB^G101S^ and *Cc*ParB^R104A^ variants were able to nucleate at the *parS* site (43). In contrast to the interaction in *C. crescentus* or *B. subtilis*, the association of *Sc*ParB with the *parS* site is not essential for CTP binding, as both the wild-type *Sc*ParB and the ScParB^HTH^ variant, which is defective in *parS* interaction, exhibit similar strengths in nucleotide binding. However, when *Sc*ParB was bound to the *parS* site, its affinity for CTP increased, suggesting that *Sc*ParB was immediately charged with CTP after the association with DNA. These findings align with those of hydrogen-deuterium exchange (HDX) studies on *Mx*ParB, which demonstrated that structural changes occurring after *Mx*ParB nucleation at the *parS* site can be transmitted across the protein, affecting the GERR motif within the NTD (12). The NTD contributes to secondary ParB dimerisation, known as N-gate closure, which is stimulated by CTP, particularly in the presence of the *parS* motif. The CTP-dependent intermolecular association of NTDs is crucial for the transformation of the ParB dimer into a clamp-like structure that encircles the *parS* site (14), facilitating the release of the ParB dimer from the *parS* site and its subsequent sliding on DNA. Consistent with these findings, only CTP-charged *Sc*ParB effectively accumulated on DNA (Figure 5E).

It also remains unclear whether ParB can bind to *parS* as a dimer or if individual ParB protomers can independently interact with the halves of the *parS* site. In our studies, we found that the G140S substitution within the CTP-binding motif also disrupted the ability of *Sc*ParB to dimerise, while its affinity for the *parS* site was unaffected. These findings suggest that *Sc*ParB can associate with the *parS* site as a monomer. Further support for this conclusion comes from studies on *B. subtilis,* which demonstrated that the *Bs*ParB dimer fails to bind both *parS* halves simultaneously because of a steric clash (14). Additionally, in *P. aeruginosa, Pa*ParB has been reported to bind 8-bp *parS*-like sequences, referred to as “half *parS”* sequences, which are hypothesised to accommodate a single *Pa*ParB protomer (3, 44). Although these findings do not entirely exclude the possibility that *Sc*ParB binds to *parS* (or half *parS*) as a dimer, with only one protomer directly interacting with DNA, they strongly suggest that the interaction between the *parS* site and a single *Sc*ParB protomer is feasible.

While CTP binding is crucial for the spread of ParB, the rate of CTP hydrolysis determines the extent to which the ParB protein can migrate beyond the *parS* site. We observed that the CTP-*Sc*ParB complex exhibited low CTP hydrolase activity when not associated with the *parS* site (10). This feature appears to be conserved among ParB homologues, serving as an autoinhibitory mechanism that prevents unnecessary CTP utilisation by ParB when the protein is not bound to DNA to fulfil its role in segrosome assembly (14). CTP hydrolysis triggers the opening of the ParB‒DNA complex, allowing ParB to be recycled in the cytoplasm (Figure 5F). Intriguingly, all the ParB homologues are relatively weak CTPases, even in the presence of *parS*-containing DNA. In *S. coelicolor,* approximately 1.2 CTP molecules are hydrolysed by *Sc*ParB per minute, which is a hydrolysis rate comparable to that of other ParB homologues (12, 14). On the other hand, the absence of CTP hydrolysis expands the chromosome region occupied by *Bs*ParB (14). Thus, CTP hydrolysis limits the sliding time of ParB on DNA, determining how far from the *parS* site ParB can migrate.

The results of our *in vitro* studies on CTP-dependent *Sc*ParB accumulation on DNA are in agreement with the observation that the complex formed *in vivo* by wild-type *Sc*ParB is significantly larger (in terms of volume) and exhibits more intense fluorescence than those formed by all the GERR-substituted *Sc*ParB-EGFP variants. Although our findings confirm the proposed model that DNA and CTP binding are essential for large segrosome formation in *S. coelicolor,* it remains unclear why the *Sc*ParB^R142A^-EGFP variant retained, at least partially, the capacity for partition complex formation. The possibility that the increased dimerisation strength observed for *Sc*ParB^R142A^-EGFP partially complements the lack of CTP-dependent accumulation on DNA cannot be excluded. On the other hand, our SIM analyses revealed that the *Sc*ParB^HTH^-EGFP variant also forms complexes with larger volume and intensity than the GERR-substituted variants. Recently, it has been shown that phase‒phase liquid separation is a feature of ParB (16, 17). *In vitro, C. glutamicum ParB* (*Cg*ParB) is able to separate into liquid-like droplets, and the phase separation is stimulated by CTP and *parS* binding. The *Cg*ParB^R175A^ variant, which carries a mutation within the Arg patch, was reported to be defective in condensation. Whereas the condensate volume of the wild-type *Cg*ParB strain reached 0.05 µm^3^, in the strain producing *Cg*ParB^R175A^, the *foci*, although still detectable, were significantly smaller than those in the wild-type strain (16). Thus, we speculate that the appearance of ParB^HTH^-EGFP *foci* in the SIM experiment may be explained by the phase separation of *Sc*ParB not bound to DNA. It was suggested that phase separation of *Cg*ParB is promoted by CTP binding in a highly crowded environment even in the absence of binding to *parS* (16).

Finally, our studies revealed that substitution within the GERR and HTH motifs affects sporogenic development. Intriguingly, the impact of particular *Sc*ParB variants on chromosome segregation, hyphal growth, and sporulation was variable and dependent on the introduced modification. Intriguingly, the *Sc*ParB^R142A^-EGFP variant, which was defective in CTP binding and hydrolysis, presented very few chromosome segregation defects, which were comparable to those of the wild-type strain. For the strains in which *Sc*ParB-EGFP *foci* were not detected, the percentage of anucleate spores increased from 7.6% (*Sc*ParB^HTH^-EGFP and *Sc*ParB^G140S^-EGFP variants) to 12.4% (*Sc*Par^G140S;R142A^-EGFP) but was still lower than that quantified for the *parB* deletion strain (17.4%) (29). This observation indicates a more pleiotropic role of the *Sc*ParB protein in chromosome partitioning, as the *Sc*ParB variants in which segrosome formation was abolished still retained some of their cellular functions.

The role of *Sc*ParB as a regulatory protein involved in sporogenic development has been reported previously. In *S. venezuelae,* the lack of *Sv*ParB or SvParA proteins resulted in slower or accelerated tip extension, respectively (23). Intriguingly, hyphal elongation is not associated with the ability of *Sc*ParB to bind to DNA or form a segrosome since the production of the *Sc*ParB^HTH^-EGFP variant resulted in a similar sporogenic cell length as that observed in the wild-type strain. On the other hand, the GERR substitutions affected the cell length but in a variable manner. While the G140S substitution resulted in shorter sporogenic compartments, similar to *parB* deletion in *S. venezuelae* (23), the G140R;R142A double amino acid substitution led to a significant increase in hyphal length. This could be explained by the impact of the introduced mutations on ParA activity. However, BTH analysis indicated that the modifications introduced in *Sc*ParB did not affect its interaction with *Sc*ParA.

In summary, our findings show that *S. coelicolor* ParB binds and hydrolyses CTP, which promotes segrosome complex assembly. However, the discovery that the conserved GERR motif is essential for CTP binding indicates its broader, unexplored role in the regulation of *S. coelicolor* sporogenic hyphal growth.

## ACKNOWLEDGEMENTS

This work was financially supported by grants from the Polish National Science Center HARMONIA grant 2016/22/M/NZ1/00122 (to M.J.S.) and OPUS grant 2023/49/B/NZ1/00781 (to M.J.S.). We thank Aleksander Czogalla and Daniel Krowarsch for their kind support in performing the BLI and CD analyses.

