## Supplementary Figures for "Significance of the CTP-binding motif for the interactions of *S. coelicolor* ParB with DNA, chromosome segregation, and sporogenic hyphal growth"

```
Cce       MESVVVGEPGMSEGRRGLGRGLSALLGEVDAAPAQ-----------APG-----------   38
Sco       ----------MSERRRGLGRGLGALIPNAPTEKSVASAALGSAASAAPGAMPLLPNERGV   50
Bsu       -------------MAKGLGKGINALF----------------------------------   13
                         :***:*:.**:

Cce       -----------------------------EQLGGSREAPIEILQRNPDQPRRTFREEDLE   69
Sco       AAAKVATLQHVSRETEELTAPQGVEGLRPPMGAHFAEVPLDAITPNPKQPRKDFDDDALA   110
Bsu       --------NQ-----------------VDLSEETVEEIKIADLRPNPYQPRKHFDDEALA   48
                                                *  :  :  ** ***: * :: *

Cce       DLSNSIREKGVLQPILVRPSPDTAGEYQIVAGERRWRAAQRAGLKTVPIMVRELDDLAVL   129
Sco       ELVTSIREVGLLQPVVVRPT--EPGRYELIMGERRFRACRELELDAIPAIVRATEDEKLL   168
Bsu       ELKESVLQHGILQPLIVRKS--LK-GYDIVAGERRFRAAKLAGLDTVPAIVRELSEALMR   105
          :*  *: : *:***::** :      *::: ****:**.:   *.::* :**  .:  :

Cce       EIGIIENVQRADLNVLEEALSYKVLMEKFERTQENIAQTIGKSRSHVANTMRLLALPDEV   189
Sco       LDALLENLHRAQLNPLEEAFAYDQLLKDFNCTHDQLADRIGRSRPQVSNTLRLLKLSPKV   228
Bsu       EIALLENLQREDLSPLEEAQAYDSLLKHLDLTQEQLAKRLGKSRPHIANHLRLLTLPENI   165
            .::**::* :*. **** :*. *::.:: *::::*. :*:** :::* :*** *  ::

Cce       QSYLVSGELTAGHARAIAAAADPV---ALAKQIIEGGLSVRETEALARKAPNLSA--GKS   244
Sco       QNRVAAGVLSAGHARALLSVDDPEEQDRLAHRIVAEGLSVRSVEEIVTLMGSRPQKPQRA   288
Bsu       QQLIAEGTLSMGHGRTLLGLKNKNKLEPLVQKVIAEQLNVRQLEQLIQQLNQNVPRETKK   225
          *. :. * *: **.*:: .  :      *.::::   *.**. * :     .      :

Cce       KGGRPPRVKDTDTQALESDLSSVLGLDVSIDHRGSTGTLTITYATLEQLDDLCNRLTRGI   304
Sco       KGPRAGSLVSPALSDLATRLSDRFETRVKVDLGQKKGKITVEFASMDDLERILGSLAPGE   348
Bsu       KEPVKD----AVLKERESYLQNYFGTTVNIKRQKKKGKIEIEFFSNEDLDRILELLSERE   281
          *            .   : *.. :   *.:.   ..*.: : : : ::*: :   *:

Cce       --------------------   304
Sco       GPVLQKGLLEGEDEDGDAES   368
Bsu       S-------------------   282
```

**Supplementary Figure 1. Alignment of ParB homologues.** The amino acid sequences of the ParB homologues were retrieved from the UniProtKB database. The accession numbers for each protein are as follows: *C. crescentus* (B8GW30, *Cce*), *B. subtilis* (P26497, *Bsu*), and *S. coelicolor* (Q9RFN2, *Sco*). The ParA-binding motif is highlighted in red, the CTP-binding motif in blue, and the helix-turn-helix (HTH) motif in green. The amino acids substituted in the *Sc*ParB variants described in this study are indicated by a red line.

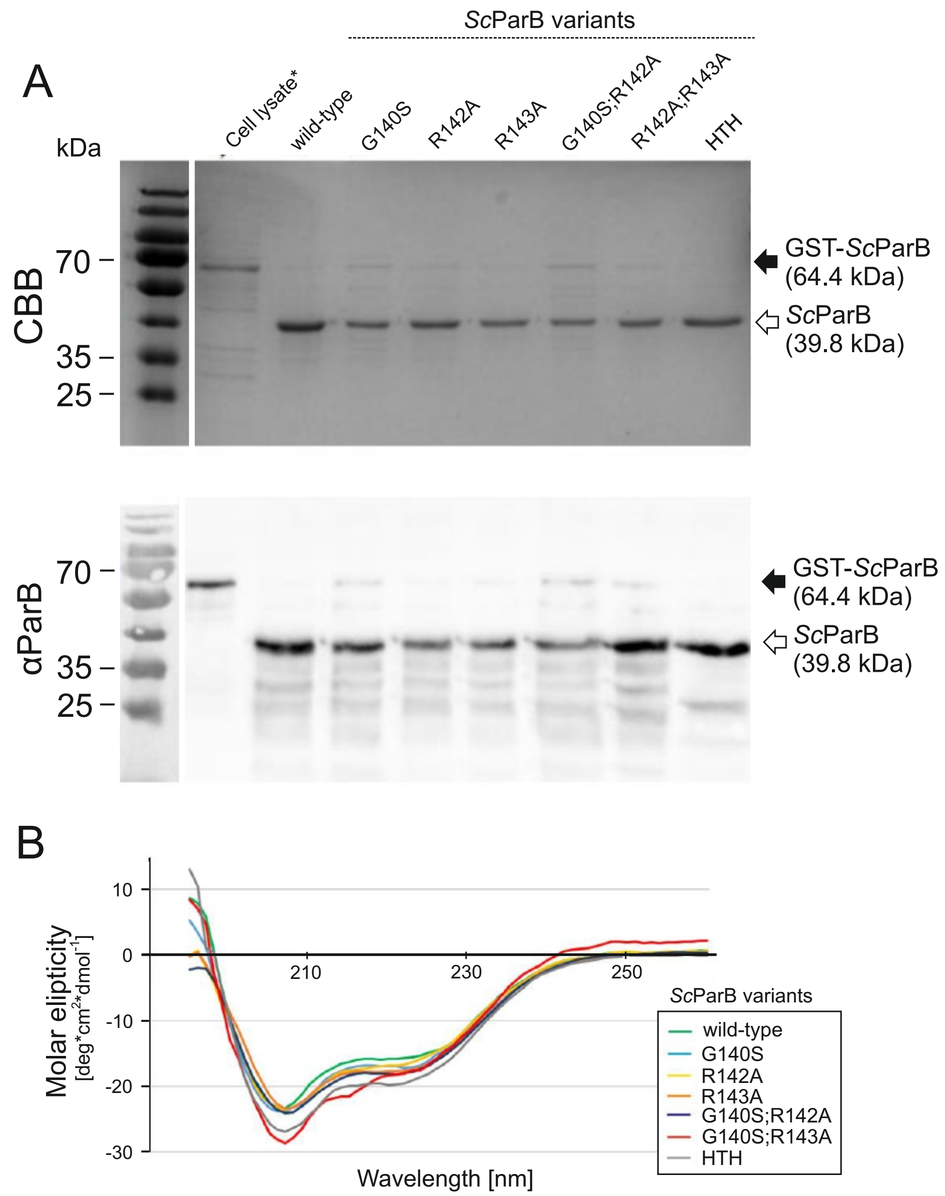


**Supplementary Figure 2. Purification of *Sc*ParB variants. (A)** Analysis of purified *Sc*ParB variants (wild-type, G140S, R142A, R143A, G140S;R142A, R142A;R143A, and HTH) using SDS–PAGE, followed by staining with Coomassie Brilliant Blue (CBB) or Western blotting with anti-ParB serum (αParB). An example of a cell lysate obtained from IPTG-induced *E. coli* BL21 (DE) pLysS cells producing wild-type GST-*Sc*ParB (64.4 kDa, black arrow) is shown (cell lysate). The purified recombinant *Sc*ParB variants after cleavage with the PreScission protease (39.8 kDa, white arrow) are indicated according to the amino acid substitutions. The 25, 35 and 70 kDa bands of the protein molecular weight ladder are marked on the left. (**B**) Normalised circular dichroism (CD) spectra of *Sc*ParB variants (wild-type, G140S, R142A, R143A, G140S;R142A, R142A;R143A, and HTH). The CD spectra were recorded in triplicate, and the average molar ellipticity [deg * $cm^2$ *$dmol^{-1}$] was plotted against the wavelength [nm].

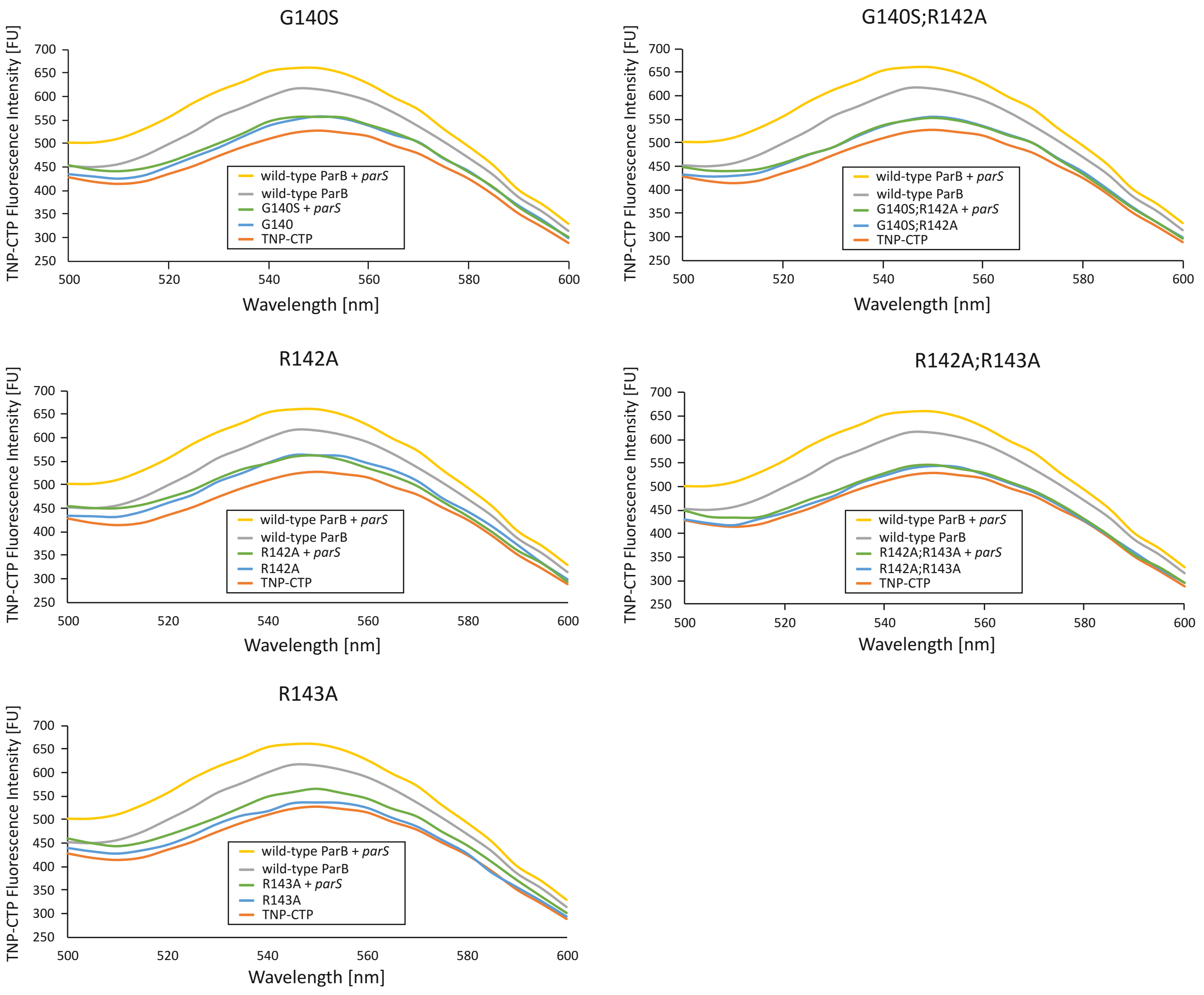


**Supplementary Figure 3. TNP-CTP binding by GERR-substituted *Sc*ParB variants.** The fluorescence intensity ([FU] spectra of 5 µM TNP-CTP measured after binding to specific *Sc*ParB variants at a concentration of 1 µM (G140S, R142A, R143A, G140S;R142A, R142A;R143A). Each analysis was conducted in the presence (green) or absence (blue) of a 34-bp *parS*-containing DNA fragment (2 µM) and compared with spectra recorded for the wild-type *Sc*ParB (yellow and grey, respectively, in the presence or absence of DNA) or a 5 µM TNP-CTP sample with no protein added (orange).

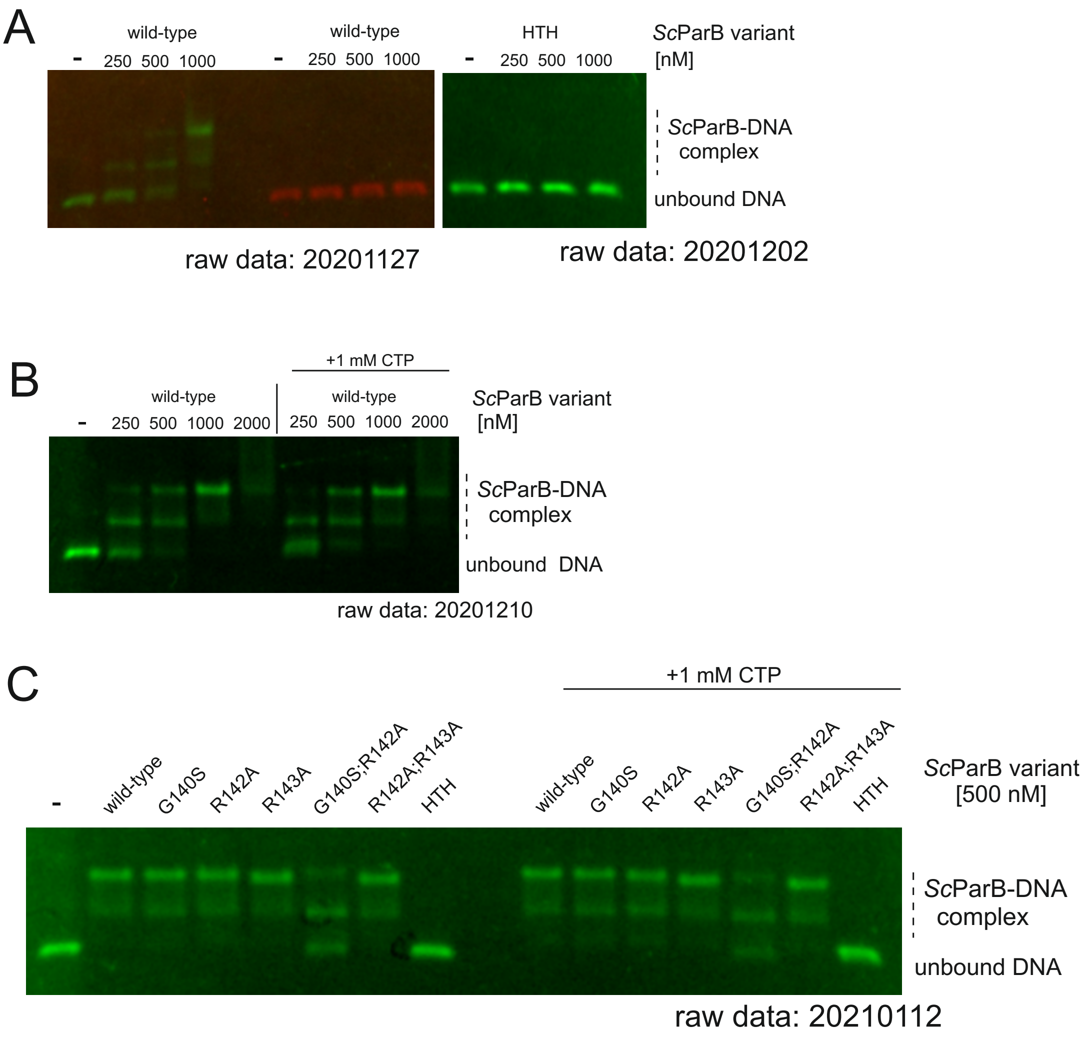


**Supplementary Figure 4. Impact of CTP on *Sc*ParB-parS binding *in vitro*** (**A**) *Left*: Analysis of recombinant wild-type *Sc*ParB (250–1000 nM) binding to a 500-bp Cy5-labelled (green) linear DNA fragment (10 µM) containing two *parS* sites or a Cy3-labelled (red) DNA fragment with two scrambled *parS* sites. *Right*: Analysis of the binding of the non-DNA-interacting *Sc*ParB variant (HTH) (250–1000 nM) to the 500-bp Cy5-labelled linear (green) DNA fragment (10 µM) containing two *parS* sites. (**B**) Analysis of recombinant wild-type *Sc*ParB (250–1000 nM) binding to the 500-bp Cy5-labelled (green) linear DNA fragment (10 µM) containing two *parS* sites in the presence (right) or absence (left) of 1 mM CTP. (**C**) Analysis of recombinant *Sc*ParB variants (wild-type, G140S, R142A, R143A, G140S;R142A, R142A;R143A, and HTH) at 500 nM in the 500-bp Cy5-labelled (green) linear DNA fragment (10 µM) containing two *parS* sites in the presence (right) or absence (left) of 1 mM CTP. The unbound DNA and *Sc*ParB-parS complexes are indicated on the right.

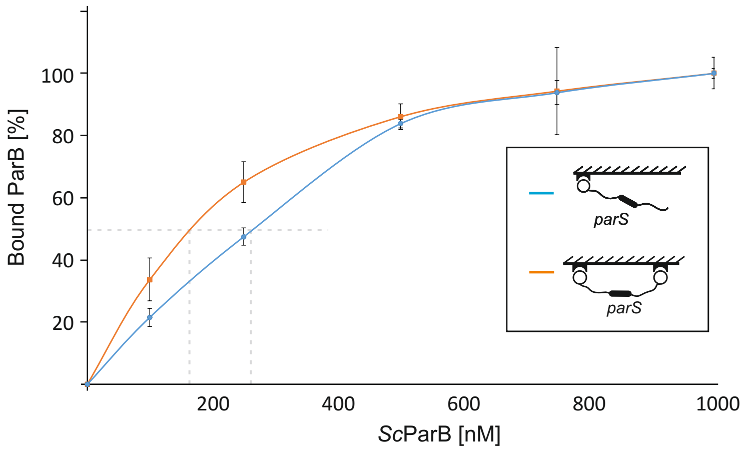


**Supplementary Figure 5. Quantification of the dissociation constant ($K_d$) of the *Sc*ParB-*parS* complex.** The Kd values were calculated on the basis of BLI measurements conducted for one- (blue) or two-end (orange) immobilised *parS*-containing DNA at the steady state of protein complex formation. The percentage of bound DNA was plotted against the *Sc*ParB concentration.

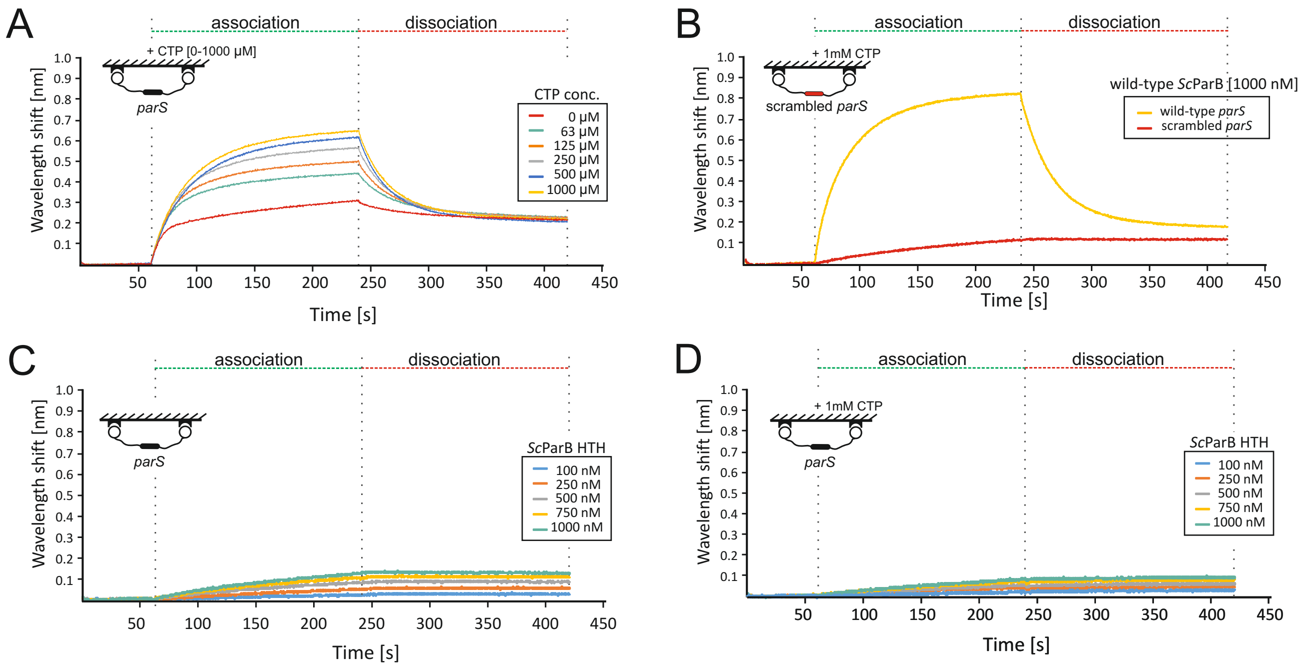


**Supplementary Figure 6. Analysis of *Sc*ParB interactions with a two-end biotin-immobilised 300-bp DNA fragment using BLI. (A)** Binding of wild-type *Sc*ParB (1000 nM) to a DNA fragment containing a wild-type *parS* site (yellow) or a scrambled *parS* site in the presence of 1 mM CTP. (**B**) Binding of wild-type *Sc*ParB (500 nM) to a DNA fragment containing a *parS* site conducted in the presence of a broad range of CTP concentrations (63–1000 µM). (**C**) Binding of the non-DNA-interacting *Sc*ParB$^{HTH}$ variant to a DNA fragment containing a wild-type *parS* site over a broad range of protein concentrations (0–1000 nM). (**D**) Binding of the non-DNA-interacting *Sc*ParB$^{HTH}$ variant to a DNA fragment containing a wild-type *parS* at a broad range of protein concentrations (0–1000 nM) in the presence of 1 mM CTP. The association and dissociation steps are indicated by green and red dotted lines, respectively.

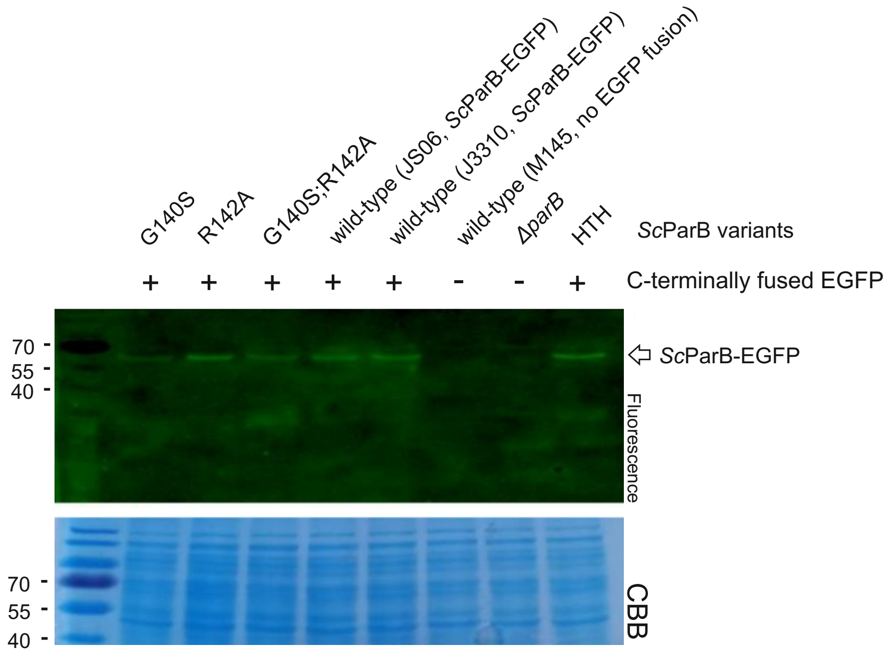


**Supplementary Figure 7. Analysis of *Sc*ParB-EGFP variant production in *S. coelicolor*.** The fluorescence signal detected in cell lysates obtained from *S. coelicolor* strains producing ScParB-EGFP variants (wild-type, G140S, R142A, G140S;R142A, and HTH). Cell lysates obtained from the wild-type (without EGFP fusion) and *parB* deletion (Δ*parB*) strains were used as controls. The loading control, represented by CBB-stained acrylamide gel (CBB), is displayed below. The 40, 55 and 70 kDa bands of the protein molecular weight ladder are indicated on the left.

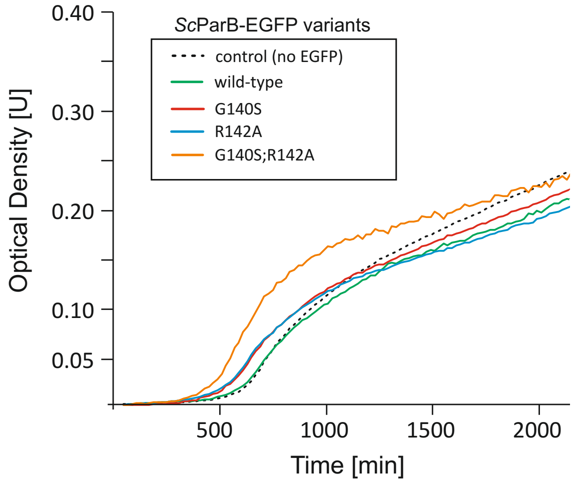


**Supplementary Figure 8. Growth of *S. coelicolor* strains producing *Sc*ParB-EGFP variants.** The growth was analysed in liquid 79 medium: wild-type [green], G140S [red], R142A [blue], G140S;R142A [orange]. The wild-type *S. coelicolor* strain, which does not produce EGFP-fused *Sc*ParB, served as a control (black dotted line).
