## Supplementary Materials and Methods for "Significance of the CTP-binding motif for the interactions of *S. coelicolor* ParB with DNA, chromosome segregation, and sporogenic hyphal growth"

**Growth rate analysis**

To obtain *S. coelicolor* growth curves, we utilised the Bioscreen C instrument (Growth Curves, US). *S. coelicolor* cell cultures were established in sterile honeycomb microplates (Growth Curves, US) containing 300 µl of liquid 79 medium inoculated with 6 × $10^3$ colony-forming units (CFUs). Data collection and analysis were conducted according to the general protocol previously described for *S. venezuelae* (36).

**Biolayer interferometry (BLI)**

To analyse the *Sc*ParB-DNA interactions, 300-bp DNA fragments containing either a single wild-type or scrambled *parS* site were amplified by PCR using the BLI_*parS*300_Fw and BLI_*parS*300_Rv oligonucleotides. The DNA templates used for the PCR were the pUC19_A7 plasmid (containing the wild-type *parS* sequence) and the pUC19_B2 plasmid (containing the scrambled *parS* sequence). To immobilise the DNA fragments, a 5'-biotin label was added to either only the forward (one-site biotinylated DNA) oligonucleotide or both the forward and reverse (two-site biotinylated DNA) custom-synthesised oligonucleotides.

BLI experiments were conducted using an Octet K2 system equipped with Octet High Precision Streptavidin (SAX) biosensors (Sartorius, US). All the experiments were performed at 30 °C. First, the streptavidin-coated biosensors were hydrated in BLI_A buffer (100 mM Tris-HCl (pH 8.0), 100 mM NaCl, 1 mM $MgCl_2$, 0,005% Tween 20) for at least 10 minutes at room temperature. Following hydration, 150 ng of one- or two-site biotinylated double-stranded DNA (dsDNA) was immobilised on the biosensors in BLI_A buffer. The DNA-coated biosensors were then washed sequentially with BLI_A buffer for 60 seconds, BLI_B buffer (100 mM Tris-HCl (pH 8.0), 1 M NaCl, 1 mM $MgCl_2$, 0.005% Tween-20, 0.1% SDS) for 300 seconds, and neutralised with BLI_A buffer for 300 seconds. Prior to each analysis, the DNA-coated biosensors were calibrated with BLI_A buffer for 60 seconds. For the measurements, *Sc*ParB variants were diluted in BLI_A buffer to final concentrations ranging from 100 to 1000 nM. The association and dissociation of the *Sc*ParB-DNA complex were monitored for 180 seconds at each step. Following each association/dissociation cycle, the biosensors were regenerated in BLI_B buffer for 300 seconds, neutralised with BLI_A buffer for 300 seconds, and then reused.

For studies examining nucleotide-dependent *Sc*ParB accumulation on DNA, *Sc*ParB (or its variant) was diluted to a concentration of 750 nM in BLI_A buffer supplemented with 1 mM nucleotide-5' triphosphates (NTPs: CTP, dCTP, TTP, GTP, or ATP). The association and dissociation steps were then recorded as described above.

**Supplementary Table 1. *Escherichia coli* strains used in the study.**

| Strain name | Relevant genotype and characteristics | Source |
|---|---|---|
| DH5α | *F-, Φ80dlacZΔM15, recA1, endA1, gyrA96, thi-E1, hsdR17, (rk-,mk+), supE44, relA1, deoR, Δ(lacZYA-argF)U169* | Laboratory stock |
| ET12567/pUZ8002 | *dam, dcm, hsdS,* CmR, TetR, pUZ8002*: tra,* KanR*, RP4 23;* | (32) |
| BL21 (DE3) pLysS | F−, *omp*T, *hsd*SB (rB−, mB−), *dcm*, *gal*, λ(DE3), pLysS, CmR | Promega (US) |
| BW25113/pIJ790 | K12 derivative; *araBAD,rhaBAD* λ-Red (*gam bet exo) cat araC rep101*(Ts) | (33) |
| BTH101 | *F-, cya-99, araD139, galE15, galK16, rpsL1* (StrR*), hsdR2, mcrA1, mcrB1* | (34) |

**Supplementary Table 2. *Streptomyces coelicolor* strains used in the study.**

| Strain | Relevant genotype and characteristics | Source |
|---|---|---|
| **M145 (wild-type)** | *S. coelicolor* M145, SCP1-, SCP2- | (35) |
| **J3303 (*ΔparB*)** | M145 *ΔparB::accIV* | This study |

| | | |
|---|---|---|
| **J3316 (*Sc*ParB$^{HTH}$-EGFP)** | M145 *parB(IGR(207-209)TYE)-egfp* | (33) |
| **JS01 (*Sc*ParB$^{G140S}$-EGFP)** | M145 *parB(G140S)-egfp* | This study |
| **JS02 (*Sc*ParB$^{R142A}$-EGFP)** | M145 *parB(R142A)-egfp* | This study |
| **JS04 (*Sc*ParB$^{G140S;R142A}$-EGFP)** | M145 *parB(G140S;R142A)-egfp* | This study |
| **JS06 (*Sc*ParB-EGFP)** | M145 *parB-egfp* | This study |

**Supplementary Table 3. Oligonucleotides used in the study.**

| **Oligonucleotide** | **Sequence** |
|---|---|
| ScParB_inside_Fw | GTTTCACGTGAAACCGAAG |
| ScParB_inside_Rv | GAAGAAGCTTCTCGTCCTC |
| ParB_EcoRI_Rv | GGGAATTCTCAGGACTCGGCGTCCCC |
| ParB_XbaI_Fw | GTTCTAGACCCAGTGAGTGAGCGACGGAGGGGG |
| ParB_KpnI_Rv | ATGGTACCATCATATGATTCAGGACTCGGCGTCCCC |
| EMSA_*parS*_Fw [1)] | CCGAGCTCGAATTCACTAGTGA TTCCG |
| EMSA_2x_*parS*_Rv | GTCGCGGCAGCCCCTCGC |
| BLI_*parS*300_Fw [2)] | GGTAGGTTATCCACGTGTTACTC |
| BLI_*parS*300_Rv [3)] | GAACCAGTGAGGCCTGGTCTTC |
| *parS*_34pz_Fw | GTGCATCGTGTTTCACGTGAAACGTCGCTCACTG |
| *parS*_34pz_Rv | CAGTGAGCGACGTTTCACGTGAAACACGATGCAC |
| mut*parS*_34pz_Fw | GTGCATCGTGTTTCTAGGGAAACGTCGCTCACTG |
| mut*parS*_34pz_Rv | CAGTGAGCGACGTTTCCCTAGAAACACGATGCAC |

[1)] The oligonucleotide was 5’-conjugated with cyanine 3 (Cy3) or cyanine 5 (Cy5) depending on the amplified DNA (see the EMSA experiment description).

[2)] The oligonucleotide was 5’-biotinylated

[3)] The oligonucleotide was 5’-biotinylated only for amplification of DNA biotinylated at both ends (see the BLI experiment description).

**Supplementary Table 4. Cosmids and plasmids used in this study**

| **Plasmid** | **Relevant genotype or characteristics** | **Source** |
|---|---|---|
| **H24_*parB* HTH-*egfp*** | SuperCos based cosmid containing fragment of *S. coelicolor* chromosome with *parB* gene fused to e*gfp* and modified with *SnaBI* restriction site disrupting nucleotide sequence within the helix-turn-helix HTH motif | (27) |
| **pGEX-6P-1_*parB*** | pGEX-6P-1 derivative encoding the *S. coelicolor parB* gene with in frame 5'-fusion with GST-encoding sequence | (26) |
| **pGEX-6P-1_*parB*(G140S)** | pGEX-6P-1_*parB* derivative, guanine(418)-to-adenine substitution in the *parB* gene | This study |
| **pGEX-6P-1_*parB*(R142A)** | pGEX-6P-1_*parB* derivative, cytosine(424)-to-guanine and guanine(425)-to-cytosine substitutions in the *parB* gene | This study |
| **pGEX-6P-1_*parB*(R143)** | pGEX-6P-1_*parB* derivative, cytosine(427)-to-guanine and guanine(428)-to-cytosine substitutions in the *parB* gene | This study |
| **pGEX-6P-1_*parB*(G140S;R142A)** | pGEX-6P-1_*parB* derivative, guanine(418)-to-adenine, cytosine(424)-to-guanine and guanine(425)-to-cytosine substitutions in the *parB* gene | This study |
| **pGEX-6P-1_*parB*(R142A;R143A)** | pGEX-6P-1_*parB* derivative, cytosine(424)-to-guanine and guanine(425)-to-cytosine, cytosine(427)-to-guanine, guanine(428)-to-cytosine substitutions in the *parB* gene | This study |
| **pGEX-6P-1_*parB*(HTH)** | pGEX-6P-1_*parB* derivative, encoding the *S. coelicolor parB* gene with *SnaBI* restriction site disrupting nucleotide sequence within the helix-turn-helix HTH motif | This study |
| **pUC19_A7** | pUC19 derivative containing 500 bp fragment of *S. coelicolor parAB* operon upstream region containing two wild-type *parS* sites | (36) |
| **pUC19_B2** | pUC19 derivative containing 500 bp fragment of *S. coelicolor parAB* operon upstream region containing two scrambled *parS* sites | (36) |
| **pUT18C** | pUC19 derivative encoding a fragment (T18 domain) of adenylate cyclase(Cya) from *Bordetella pertussis* | (34) |
| **pKT25** | pSU40 derivative encoding a fragment (T18 domain) of adenylate cyclase (Cya) from *Bordetella pertussis* | (34) |
| **pUT18C_*parB*** | pUT18C derivative encoding the *S. coelicolor* wild-type *parB* gene | This study |
| **pUT18C_*parB*(140S)** | pUT18C derivative, encoding the *S. coelicolor parB* gene with guanine(418)-to-adenine substitution | This study |

| | | |
|---|---|---|
| **pUT18C_*parB*(142A)** | pUT18C derivative, encoding the *S. coelicolor parB* gene with cytosine(424)-to-guanine and guanine(425)-to-cytosine substitutions | This study |
| **pUT18C_*parB*(143A)** | pUT18C derivative, encoding the *S. coelicolor parB* gene with cytosine(427)-to-guanine and guanine(428)-to-cytosine | This study |
| **pUT18C_*parB*(G140S;R142A)** | pUT18C derivative, encoding the *S. coelicolor parB* gene with guanine(418)-to-adenine, cytosine(424)-to-guanine and guanine(425)-to-cytosine substitutions | This study |
| **pUT18C_*parB*(R142A;R143A)** | pUT18C derivative, encoding the *S. coelicolor parB* gene with cytosine(424)-to-guanine and guanine(425)-to-cytosine, cytosine(427)-to-guanine, guanine(428)-to-cytosine substitutions | This study |
| **pUT18C_*parB*(HTH)** | pUT18C derivative, encoding the *S. coelicolor parB* gene with *SnaBI* restriction site disrupting nucleotide sequence within the helix-turn-helix HTH motif | This study |
| **pUT18C_parA** | pUT18C derivative encoding the *S. coelicolor* wild-type *parA* gene | (29) |
| **pKT25_*parB*** | pKT25 derivative encoding the *S. coelicolor* wild-type *parB* gene | This study |
| **pKT25_*parB*(140S)** | pKT25 derivative, encoding the *S. coelicolor parB* gene with guanine(418)-to-adenine substitution | This study |
| **pKT25_*parB*(142A)** | pKT25 derivative, encoding the *S. coelicolor parB* gene with cytosine(424)-to-guanine and guanine(425)-to-cytosine substitutions | This study |
| **pKT25_*parB*(143A)** | pKT25 derivative, encoding the *S. coelicolor parB* gene with cytosine(427)-to-guanine and guanine(428)-to-cytosine | This study |
| **pKT25_*parB*(G140S;R142A)** | pKT25 derivative, encoding the *S. coelicolor parB* gene with guanine(418)-to-adenine, cytosine(424)-to-guanine and guanine(425)-to-cytosine substitutions | This study |
| **pKT25_*parB*(R142A;R143A)** | pKT25 derivative, encoding the *S. coelicolor parB* gene with cytosine(424)-to-guanine and guanine(425)-to-cytosine, cytosine(427)-to-guanine, guanine(428)-to-cytosine substitutions | This study |
| **pKT25_*parB*(HTH)** | pKT25 derivative, encoding the *S. coelicolor parB* gene with *SnaBI* restriction site disrupting nucleotide sequence within the helix-turn-helix HTH motif | This study |
